# Hyperpolyploidization of hepatocyte initiates preneoplastic lesion formation in the liver

**DOI:** 10.1101/2020.08.23.258699

**Authors:** Heng Lin, Yen-Sung Huang, Jean-Michel Fustin, Masao Doi, Huatao Chen, Hui-Huang Lai, Shu-Hui Lin, Yen-Lurk Lee, Pei-Chih King, Hsien-San Hou, Hao-Wen Chen, Pei-Yun Young, Hsu-Wen Chao

## Abstract

Hepatocellular carcinoma (HCC) is the most predominant primary malignancy in the liver. Genotoxic and genetic models have revealed that HCC cells are derived from hepatocytes, but where the critical region for tumor foci emergence is and how this transformation occurs are still unclear. Here, hyperpolyploidization of hepatocytes around the centrilobular (CL) region was demonstrated to be closely linked with the development of HCC cells after diethylnitrosamine treatment. We identified the CL region as a dominant lobule for accumulation of hyperpolyploid hepatocytes and preneoplastic tumor foci formation. We also demonstrated that upregulation of *Aurkb* plays a critical role in promoting hyperpolyploidization. Increase of AURKB phosphorylation was detected on the midbody during cytokinesis, causing abscission failure and hyperpolyploidization. Pharmacological inhibition of AURKB dramatically reduced nucleus size and tumor foci number surrounding the CL region in diethylnitrosamine-treated liver. Our work reveals an intimate molecular link between pathological hyperpolyploidy of CL hepatocytes and transformation into HCC cells.

## INTRODUCTION

Hepatocellular carcinoma (HCC) is the most predominant primary malignancy in the liver, accounting for 90% of all liver cancer cases and is characterized by poor survival rate^1,2^. HCC develops progressively, after long-term exposure to carcinogenic agents causing random genetic variations^3^. Stemness traits, normally lacking from the adult liver, have been found in human HCC, raising the stem/progenitor cell origin hypothesis of liver cancers^4,5^. Recent findings however suggested that HCC is fairly heterogenous and arises as a consequence of hepatocytes transformation during hepatocarcinogenesis^6,7^. Genetic lineage tracing approaches showed that HCC and hepatocellular adenoma (HCA) are derived from mature hepatocytes in mouse models^6,7^. Moreover, adult hepatocytes can trans-differentiate into biliary-like cells during liver cancer formation, and de-differentiate into progenitor-like cells in p53-deficient mouse liver^8,9^. The relevance of these models for human HCC is still a matter of debate. Despite increasing evidence on the origin of HCC cells, how hepatocytes transform into preneoplastic cells is still unclear.

Polyploidization has been proposed to play a critical role during tumorigenesis^10^. Most mammalian species are diploid, but polyploidy occurs in specific organs, such as in the liver during development, and has been suggested to signal the termination of differentiation or to enhance protection against stress conditions^11,12^. During postnatal development, hepatocytes progressively develop polyploidy^11,13^ via incomplete cytokinesis without contractile ring formation, which is the major mechanism for generation of polyploid hepatocytes after weaning^11^. Recently, a population of hepatocytes failing abscission has been identified following the disruption of the *Period* genes, which was shown to increase abscission failure and accelerate hepatocytes hyperpolyploidization^13^. “Physiological” hepatocyte polyploidization has been proposed as a diversity factor for liver homeostasis, and as a mechanism to restrict proliferation and promote tumor suppression^14,15,16^. The consequences of pathological hyperpolyploidy, for example when induced by chronic stress or exposure to carcinogens, are still largely unknown.

Under chronic stress, the adult liver has a remarkable ability to generate hyperpolyploid hepatocytes^17,18,19,20^. Recent study revealed that hyperpolyploid hepatocytes are associated with worse prognosis in human liver HCC, and that HCCs characterized by a low degree of differentiation and *TP53* mutations have higher levels of polyploidy^21^. The occurrence of hyperpolyploidy at early stages of tumor formation is believed to increase genomic instability and to be a pivotal step during tumorigenesis^22,23,24^. Hyperpolyploid giant cells, such as human ovarian, breast, colon, and prostate cancer cell lines, have been demonstrated to serve as a source of stemness and tumor heterogeneity through genomic reduction pathways^25^. Whether this is a shared feature of hyperpolyploid cells in different tissues is unknown.

Here, the pathological significance of genotoxin-induced hyperpolyploid hepatocytes around the centrilobular (CL) region is demonstrated. Hyperpolyploid CL hepatocytes are an oncogenerative source of preneoplastic cells through uncharacterized genome-reduction processes. Upregulation of *Aurkb* is identified to cause abscission failure and promote hyperpolyploidization of hepatocytes. Our findings show that, under treatment with diethylnitrosamine (DEN), a carcinogenic compound known to cause HCC, hepatocytes hyperpolyploidization is a crucial step for the transformation of hepatocytes into HCC cells.

## RESULTS

### DEN causes pathological hyperpolyploidization of CL-hepatocytes

To address how hepatocytes transform into HCC cells, HCC was induced with DEN, and liver tissues were examined histologically at the preneoplastic stage. DEN-injected mice were sacrificed at the indicated times after 5h BrdU-labeling (Fig. 1a). Global liver morphology showed no significant change up to three months after DEN injection, although very few small tumor nodules were sometimes found on the surface of DEN-treated liver (Fig. 1b). The hepatocytes from older control mice at p105 had larger nuclei compared to younger control mice, but DEN-treated hepatocytes still display significantly bigger cells and nuclei than age-matched control mice (Fig. 1c,d and Supplementary Fig. 1b-g), in line with previous findings^17^. Flow cytometry analysis also demonstrated that enlarged nuclei correlated with higher DNA content in DEN-treated hepatocytes (Supplementary Fig. 1d). Because the zonal distribution of hepatocytes has been proposed to be relevant to the pharmacokinetic of DEN^26^, we next investigated whether the DEN-induced hyperpolyploid hepatocytes were zone-specific. Using glutamine synthetase (GS) expression as marker for CL hepatocytes^21^, the central vein to portal vein (CV-PV) axis was subdivided into 15 parts (Supplementary Fig. 1a), and the sizes of cells and nuclei were quantified along this axis, as described previously^13^. Remarkably, only hepatocytes close to the CL and midlobular (ML) regions displayed enlarged nuclei compared to control liver (Fig. 1c,d and Supplementary Fig. 1c-g). Further examination of the effects of DEN on hepatocytes revealed these enlarged nuclei and cells became gradually evident around 2-3 months in DEN-treated liver (Fig. 1c,d and Supplementary Fig. 1f,g). In contrast, perilobular (PL) hepatocytes displayed similar nucleus sizes compared to age-matched control liver (Fig. 1c,d). To investigate whether hyperployploidization of CL hepatocytes is simply an artifact of the DEN model or a relevant feature of early-stage HCC formation caused by xenobiotics or oxidative stress, three different mouse models, using Aflatoxin B1 (AFTB1), carbon tertrachloride (CCl_4_), and 45 kcal% high-fat diet (HFD) were utilized to address this question. As shown in Supplementary Fig. 1k, hepatocytes in AFTB1- and CCl_4_-treated mice displayed enlarged nucleus size near the central vein region, compared to control group. After 90 days of HFD treatment, hepatocytes also exhibited slightly larger nucleus size than age-matched control nearby central vein region, but not in the liver with 60 days of HFD treatment. This result indicated that not only DEN, but also xenobiotics and HFD-induced oxidative stress specifically target CL and ML hepatocytes and cause hepatic hyperpolyploidization within these two regions. More generally, the metabolic sensitivity of CL and ML hepatocytes to xenobiotics is likely an important factor in HCC development.

**Figure 1.**
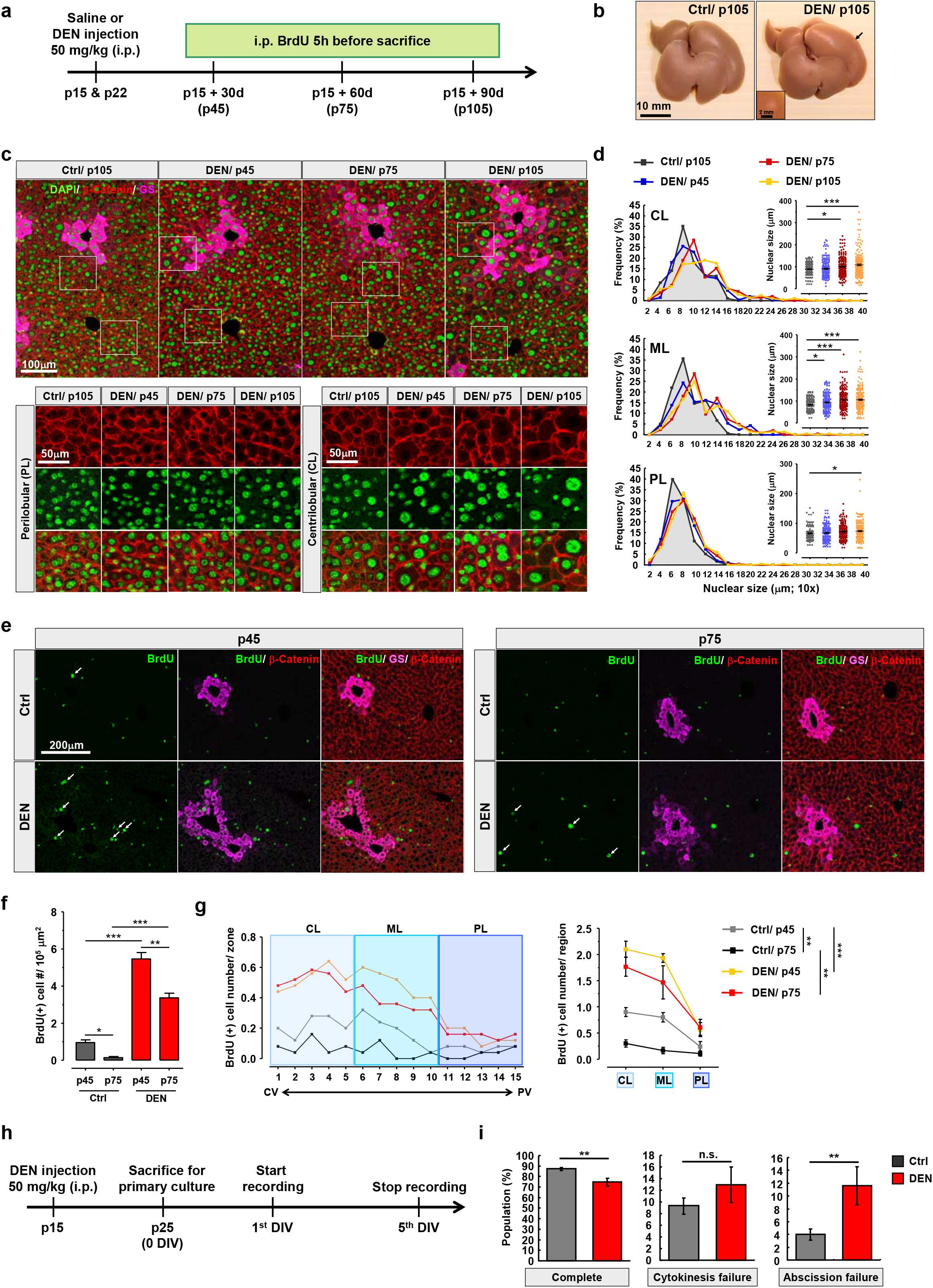
Emergence of hyperpolyploid hepatocytes within CL and ML regions of DEN-treated liver. (**a**) Schema for DEN-injection experiment: Injection of DEN (50 mg/kg) or normal saline (Ctrl) was performed at p15 and p22. Mice were injected with BrdU, 5h before sacrifice. (**b**) Example of images shows the global morphology of liver. (**c**) Representative images of liver sections from indicated mice. DEN-injected mice were sacrificed at indicated times, and control mice have the same age like mice with 3 months of DEN treatment. The higher magnification shows the detail morphology of CL and PL hepatocytes. (**d**) The nucleus size of hepatocytes along CV-PV axis of liver. Mouse number (N) = 5 mice/group, and cell number (n) = 750 cells/group. (**e**) Images of sections were prepared from p45 and p75 liver tissues with or without DEN injection, describing in Fig 1a. (**f**) The numbers of BrdU positive hepatocytes in indicated liver at p45 and p75 age. N = 5 mice/group. (**g**) Zonal distribution of BrdU positive hepatocyte along CV-PV axis. CV-PV axis is divided into 15 parts. Part 1 to 5, 6 to 10, and 11 to 15 stand for CL, ML, and PL of the liver respectively. N = 5 mice/group. (**h**) Schematic of hepatocyte primary culture for time-lapse recording: Mice are injected with DEN or normal saline at p15 and sacrificed at p25 for primary culture. 24 hours after seeding, the first day *in vitro* (DIV), hepatocytes were conducted to cell recording. (**i**) The ratio of indicated cytokinesis behaviors. Data are representative of at least three independent experiments. n = 276 and 260 dividing cells for control and DEN-treated groups respectively. Statistic: One-way ANOVA, (**d**), and (**f**); Two-way ANOVA, (**g**); Mann-Whitney nonparametric test, (**i**). Values represent the mean ± s.e.m., \**P* < 0.05, \*\**P* < 0.01, \*\*\**P* < 0.001. n.s., not significant.

The understand the cause of hyperpolyploidy, we next investigated whether cell proliferation or cytokinesis was affected by DEN. We recorded the number and distribution of BrdU-positive hepatocytes along the CV-PV axis. In control liver, the number of BrdU-positive hepatocytes progressively decreased during development, consistent with our previous finding (Fig. 1e,f and Supplementary Fig. 1h,i)^13^. In DEN-injected liver, however, the number of BrdU-positive hepatocytes increased (Fig. 1e,f). Importantly, the distribution of BrdU-positive hepatocytes was enriched around the CV, thus highly correlated with the distribution of hyperpolyploid hepatocytes (Fig. 1e,g and Supplementary Fig. 1h,i). As anticipated, BrdU-positive hepatocytes displayed enlarged nuclei, compared to age-matched control BrdU-positive hepatocytes (Supplementary Fig. 1j).

Next, primary hepatocytes isolated from liver tissues with or without DEN-treatment were cultivated to follow the progression of cytokinesis by time-lapse microscopy (Fig. 1h). Mitosis successfully proceeded with no apparent abnormality, and the respective proportions of cytokinesis failure without contractile ring formation showed no differences between the two groups (Fig. 1i and Supplementary Video 1,2). However, after DEN treatment the frequency of abscission failure was significantly increased, in parallel with a reduced ratio of complete cytokinesis (Fig. 1i and Supplementary Video 3). Interestingly, we found that bi-nucleated hepatocytes with enlarged nuclei were generated by two successive cycles with abscission failure (Supplementary Video 4), while hepatocytes with a single macronucleus were produced via abscission failure followed by a complete cytokinesis (Supplementary Video 5). Overall, these results indicated that aberrant cell proliferation coupled with abscission failure underlies DEN-induced hepatocyte hyperpolyploidization.

### Preneoplastic lesions mainly originate within the CL region

To investigate the correlation between hyperpolyploidy and the formation of preneoplastic lesions, we first asked whether hyperpolyploid hepatocytes express precancerous markers. Nuclear vacuolation, a senescence marker identified as a precancerous phenotype^27,28,29,30^, increased in DEN-injected livers (Fig. 2a and Supplementary Fig. 2a,b). Further analysis indicated that nuclear size was positively correlated with vacuolation number and area (Fig. 2b and Supplementary Fig. 2c). Importantly, nuclear vacuolation progressively increased after DEN injection (Fig. 2c), and hepatocytes with nuclear vacuolation were also highly enriched within the CL and ML regions, similar to the distribution of hyperpolyploid hepatocytes (Fig. 2d and Supplementary Fig. 2a,b). Furthermore, senescence and DNA damage were examined by Lamin B1 and γH2AX, respectively (Supplementary Fig. 2d-f). In DEN-treated liver, higher expression of γH2AX was detected specifically around CL hepatocytes and preneoplastic foci, indicating severe DNA damage in these regions (Supplementary Fig. 2d,f). In contrast with γH2AX, downregulation of Lamin B1 was observed in DEN-treated liver, which was specifically identified in hepatocytes but not in other cell types such as portal triad endothelial cells (yellow arrow). Importantly, lowest Lamin B1 expression was seen in preneoplastic cells, compared to adjacent hepatocytes (Supplementary Fig. 2e,f). Because hyperpolyploid hepatocytes with precancerous traits were mainly located within the CL and ML regions, we next sought to determine whether preneoplastic lesions (see definition in Supplementary material) also predominantly occurred in these regions. We found that microscopic foci of preneoplastic lesions were first observed two months after DEN injection, the number of preneoplastic lesions increasing thereafter (Fig. 2e). These preneoplastic lesions were not only BrdU- and Ki67-positive, but also expressed higher levels of α-fetoprotein, a precancerous marker of HCC, with higher nuclear-to-cytoplasmic ratio in area compared to normal cells. (Fig. 2f-h, and Supplementary Fig. 2g). Importantly, the distribution of preneoplastic lesions was indeed enriched around the CV, thus highly correlated with the distribution of hyperpolyploid hepatocytes (Fig. 2f,g,i and Supplementary Fig. 2g,h).

**Figure 2.**
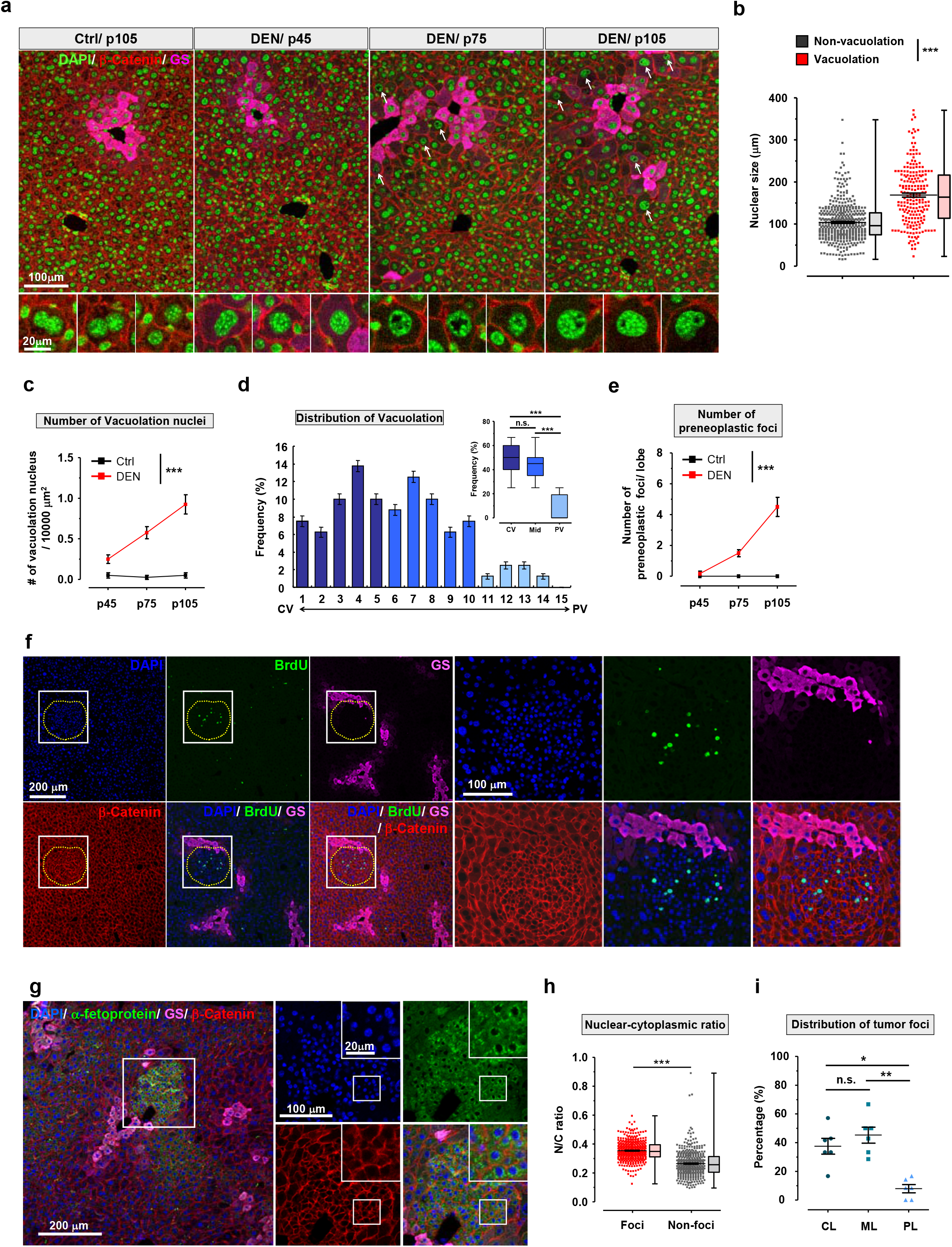
CL and ML region are the dominant areas for preneoplastic lesions generation. (**a**) Representative Images of liver sections from control (normal saline) and DEN-treated mice at indicated times. The high magnification shows the detail morphology of nuclear vacuolation in hepatocytes. (**b**) The quantitative data of nucleus size of indicated hepatocytes. N = 5 mice/group, and n = 420 and 208 nuclei from non-vacuolation and vacuolation hepatocytes respectively. (**c**) The numbers of vacuolation nuclei in control and DEN-treated liver at indicated times. N = 5 mice/group. (**d**) Bar graph displays the frequency distribution of hepatocytes with nuclear vacuolation along the CV-PV axis in DEN-treated liver. Inset shows the statistic analysis of nuclear vacuolation between CL, ML, and PL region, N = 5 mice. (**e**) The numbers of preneoplastic foci in control and DEN-treated liver at indicated times. N = 5 mice/group. (**f**) Example of images shows the distribution of preneoplastic foci nearby GS positive hepatocytes. The high magnification represents the enrichment of BrdU positive hepatocytes within preneoplastic foci. (**g**) Images display the α-fetoprotein positive signal within preneoplastic foci. The high magnification images display the detail information of fluoresce signals within preneoplastic foci. (**h**) The nuclear-cytoplasmic ratio of hepatocytes within indicated region. n > 400 hepatocytes, 6 mice/group. (**i**) The emergence frequency of preneoplastic foci within indicated liver lobule. N = 6 mice. Statistic: Student’s unpaired t-test, (**b**), and (**h**); Two-way ANOVA, (**c**), and (**e**); One-way ANOVA (**d**), and (**i**). Values represent the mean ± s.e.m., \**P* < 0.05, \*\**P* < 0.01, \*\*\**P* < 0.001. n.s., not significant.

### Preneoplastic cells are derived from CL hepatocytes and show nucleus size reduction

Surprisingly, the preneoplastic cells analyzed above were extremely small, and displayed smaller nuclei compared to cells in other areas of the liver, suggesting lower genomic content (Fig. 3a). Because reduced genomic content in human and rodent liver cancer cells has been previously demonstrated^31,32,33^, we hypothesized that preneoplastic cells originate from hyperpolyploid hepatocytes through genomic content and nucleus size reduction. If this is the case, preneoplastic cells would appear predominantly among CL hepatocytes. To test this conjecture, CL and PL hepatocytes were separately enriched by digitonin-collagenase perfusion system after one month of DEN treatment (Fig. 3b)^34,35^, when hyperpolyploid hepatocytes emerge but before preneoplastic foci formation. Digitonin caused a regularly scattered discoloration pattern on the liver surface and H&E stained sections (Fig. 3c,d). Additionally, expression of liver zone-specific markers confirmed the efficient enrichment of PL or CL hepatocytes (Fig. 3e). Importantly, microscopy and flow cytometry showed that CL hepatocytes did have bigger nuclei and higher DNA content compared to those isolated from the PL region after DEN treatment (Fig. 3f and Supplementary Fig. 3a,b).

**Figure 3.**
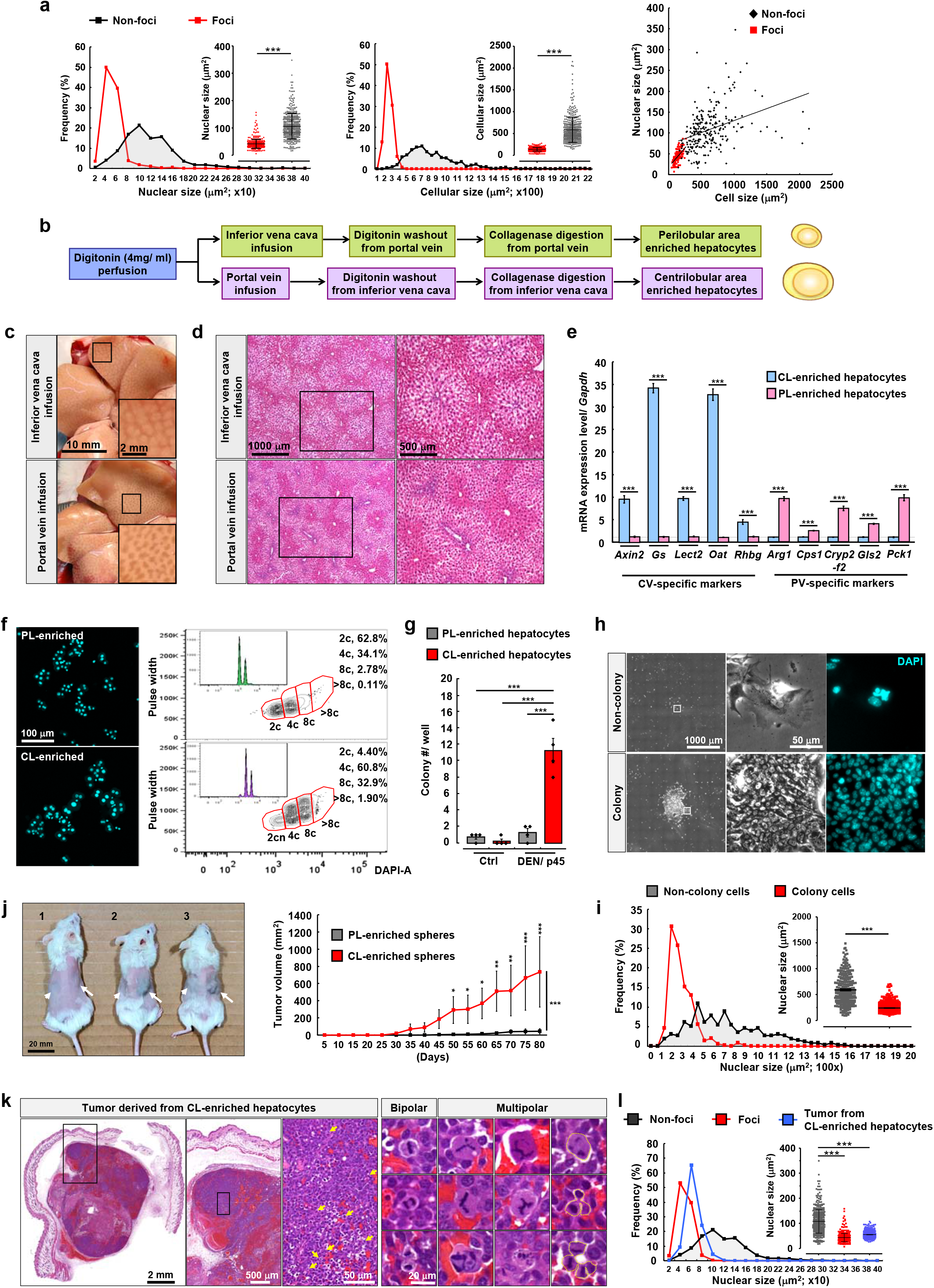
Preneoplastic cells are derived from hyperpolyploidy-enriched CL-region. (**a**) Nucleus and cell size within preneoplastic foci and non-preneoplastic region. n > 400 hepatocytes/group, N = 6 mice/group. (**b**) Schema for isolation of PL- and CL-enriched hepatocytes by digitonin-collagenase perfusion system. (**c**) Representative images display discoloration pattern on liver surface after digitonin infusion from inferior vena cava or portal vein respectively. (**d**) Representative images show the H&E staining of indicated liver histologically. Digitonin infusion caused discoloration of eosin signal. (**e**) qPCR analysis of mRNA levels of CL- and PL-specific markers. N = 3 mice/group. (**f**) The nucleus morphology of hepatocytes isolated by digitonin-collagenase infusion. The DNA contents of CL- and PL-enriched hepatocytes were analyzed by flow cytometry. (**g**) The colony numbers at 21^th^ day *in vitro* (DIV) in indicated cultured hepatocytes. (**h**) Representative images of colony and non-colony hepatocytes morphology isolated from CL-region at 21^th^ DIV. Low and high magnification show the morphology of colonies and nuclei respectively. (**i**) Nucleus size in colony and non-colony hepatocytes at 21^th^ DIV. (**j**) Visible differences in tumor size under the skin of the dorsal body surface in NSG™ mice after 10 weeks of subcutaneous transplantation. Arrow and arrow head indicate subcutaneous implantation of hepatic spheres derived from CL- and PL-enriched hepatocytes, respectively. Enlarged tumors were observed in CL-enriched hepatocytes transplanted region in j-2 and −3 mice. No tumors were detected in j-1 mouse. Line graphs show the tumor growth cure of subcutaneous transplants. N = 6 mice. (**k**) Example of images show H&E staining of tumor derived from CL-enriched hepatocytes under low and high magnification. Arrows (cyan) indicates the proliferating tumor cells at metaphase. The highest images display bipolar and multipolar dividing cells in tumor nodes. Dashed lines delineate cell borders. (**l**) Line and dot graph show the quantitative data of nucleus size within preneoplastic foci, non-preneoplastic region and tumor derived from CL-enriched hepatocytes. n > 200 cells/group. Statistic: Student’s unpaired t-test, (**a**). One-way ANOVA (**g**,**l**). Two-way ANOVA, (**j**). Values represent the mean ± s.e.m., \**P* < 0.05, \*\**P* < 0.01, \*\*\**P* < 0.001.

To examine whether CL hepatocytes displayed cancerous properties after DEN injection, colony formation assay was performed with CL or PL hepatocytes, the numbers of colonies analyzed on the 21^st^ day *in vitro* (DIV). CL hepatocytes showed higher colony formation ability compared to PL hepatocytes and to control hepatocytes isolated from the liver without DEN treatment (Fig. 3g, Supplementary Fig. 3c, and Supplementary Video 6, 7). Further analysis of the nuclei demonstrated that colony-forming cells had significantly smaller nuclei compare to non-colony cells (Fig. 3h,i). Next, tumorsphere formation assay was applied to investigate whether CV hepatocytes were more like cancer stem/progenitor cells than PL hepatocytes after DEN injection. Most hepatocytes died within 48h of seeding, but a population of spherical formations was observed 7 days after seeding. Huge tumorspheres from CL hepatocytes were easily distinguishable from aggregated cells on the 14^th^ DIV and grew further until the 21^th^ DIV (Supplementary Fig. 3d). Moreover, CL hepatocytes more frequently formed tumorspheres (diameter > 100 μm) than PL hepatocytes (Supplementary Fig. 3e), and the tumorspheres derived from CL hepatocytes were larger than those from PL hepatocytes (Supplementary Fig. 3f). To address the tumorous potency of CL and PL hepatocytes after DEN treatment *in vivo*, subcutaneous implantation of spheres derived from CL- and PL-enriched hepatocytes into the right and left flanks of NSG™ immunodeficient mice was performed, respectively. Clumps were found in the right flanks (CL-enriched hepatocytes) 30 days post-injection, became significantly larger than those observed in the left flanks at 50 days of injection, and further grew thereafter, while no clumps were detected in the left flanks (Fig. 3j). H&E staining and immunohistochemistry revealed the severe angiogenesis and high expression of the cell proliferation marker Ki-67 within the clumps, confirming the tumorigenicity of CL hepatocyte-derived spheres after DEN treatment (Fig. 3k and Supplementary Fig. 3g). To investigate whether hyperpolyploid hepatocytes are the origin of the smaller preneoplastic cells, the size of the nuclei of cells in the clump was further analyzed. During isolation of CL hepatocytes, very few cells with small nuclei were observed, but 80 days after implantation cells in the CL hepatocytes-derived clumps had significantly smaller nuclei compared to CL hepatocytes in DEN-treated liver (Fig. 3l). Interestingly, multipolar dividing cells reported by Grompe’s group as the reversion mechanism of polyploid hepatocytes were detected (around 5% incidence rate) within CL hepatocytes-derived tumors (higher magnification images of Fig. 3k)^36,37^. This observation raises the possibility that “reductive mitoses” can happen in CL hepatocyte-derived tumors and results in polyploidy reversal to produce daughter cells with lower DNA content. Together these results show that hyperpolyploid hepatocytes have high tumorigenicity and may be the origin of the small preneoplastic cells seen after DEN treatment.

### Upregulation of *Aurkb* in DEN-injected liver, NAFLD and HCC patients

Because abscission failure is involved in DEN-induced hepatocyte hyperpolyploidization, we next sought to identify the mechanisms underlying abscission failure. Recently, high-throughput screens have identified critical genes involved in cytokinesis^38,39,40^. Of these 251 cytokinesis-related genes, 185 genes were also reported in two omics studies of DEN-regulated genes in rat liver (GSE19057 and GSE63726) (Fig. 4a and Supplementary Fig. 4a)^41,42^. Only 5 out of 185, including *Anxa2*, *Aurkb*, *Cdk1*, *Rhoc*, and *S100a6*, displayed a significant upregulation (>2-fold) in both datasets (Fig. 4b and Supplementary Fig. 4b). We confirmed by qPCR that four of these candidates exhibited a significant increase in mRNA expression in the liver after one month of DEN injections (Fig. 4c), with *Aurkb* and *Cdk1* increasing over five-fold. Interestingly, *Aurkb* is a master controller of the chromosomal passenger complex, and plays a key role in regulating the NoCut pathway of abscission^43,44,45^. Upregulation of AURKB is highly correlated with abscission failure, polyploidization and tumor formation^43,44,46^. Focusing on this interesting candidate, we found that the expression and activity (by phosphorylation of Thr232, pT232) of AURKB increased in DEN-injected liver, together with Histone-H3 Ser10 phosphorylation, a direct target of AURKB (Fig. 4d).

**Figure 4.**
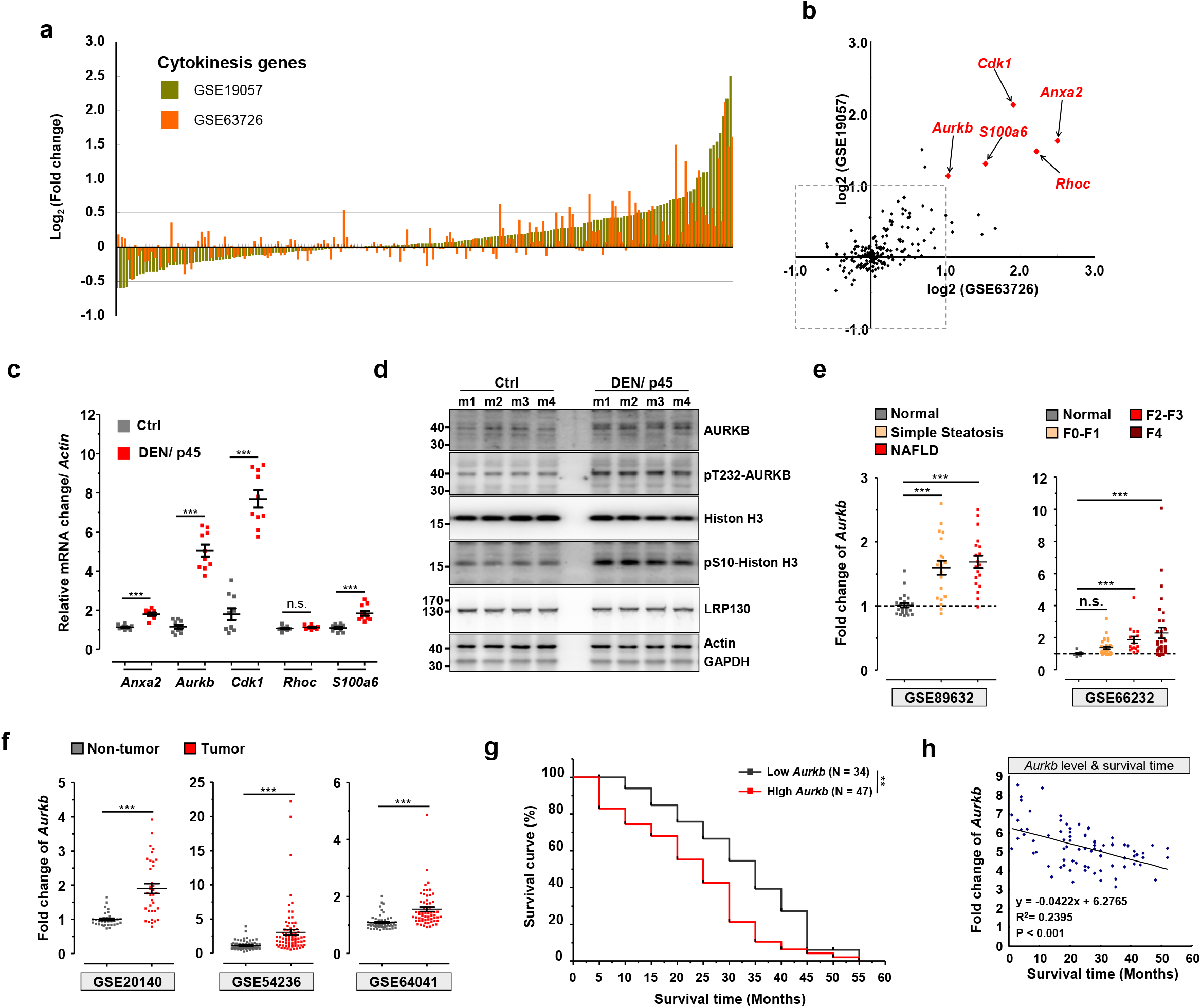
Upregulation of *Aurkb* in patients with liver diseases. (**a**) The rank order of mRNA expression levels of cytokinesis genes from GEO datasets (GSE19057 and GSE63726). (**b**) Scatter plot graph with logarithmic axes shows the top five cytokinesis genes upregulated simultaneously in two GEO datasets. (**c**) qPCR analysis of mRNA expression levels of the top five cytokinesis genes in the liver. N = 10 mice/group. (**d**) Immunoblot detection of AURKB and pT232-AURKB levels in indicated mouse liver tissues. (**e**) The expression levels of *Aurkb* from human liver biopsies in GEO datasets. Steatosis and NAFLD patients (left panel), different fibrosis stage of human livers (right panel). Patient numbers: GES89632, N = 24/20/19; GES66232, N = 10/32/16/33. (**f**) Human *Aurkb* expression in tumor and non-tumor biopsies from GEO datasets. Patient numbers: GES20140, N = 34/35; GES54236, N = 80/81; GES64041, N = 60/60. (**g**) Kaplan–Meier plots for survival curves of patients with low and high *Aurkb* expression in the liver. Patient numbers: GES54236, N = 44/37. (**h**) The linear regression analysis of the correlation between *Aurkb* expression and survival time of patients. Each point represents an individual case from a patient. Patient numbers: GES54236, N = 81. Statistic: Mann-Whitney nonparametric test, (**c**); One-way ANOVA, (**e**); Student’s unpaired t-test, (**f**); Log-rank (Mantel-Cox) Test; (**g**); Linear regression, (**h**). Values represent the mean ± s.e.m., \**P* < 0.05, \*\**P* < 0.01, \*\*\**P* < 0.001.

To evaluate whether higher expression of *Aurkb* is positively correlated with human liver diseases, we first mined GEO datasets from human samples. Patients with simple steatosis and NAFLD in the GSE89632 dataset show a significant increase of *Aurkb* expression in the liver (Fig. 4e), echoing the similar findings of pathological polyploidization in NAFLD livers^20^. Furthermore, the expression of *Aurkb* gradually increased during liver fibrosis progression in GSE66232 dataset (Fig. 4e). In support of its putative oncogenic activity, significant upregulation of *Aurkb* was found in HCC patients from at least three GEO datasets (GSE20140, GSE54236, and GSE64041) (Fig. 4f), and *Aurkb* induction was confirmed in matching pairs of HCC and normal tissues in GES64041-3 dataset (Supplementary Fig. 4c). Importantly, *Aurkb* expression showed significantly negative correlation with tumor doubling time and patient survival (Fig. 4g,h and Supplementary Fig. 4d). These results efficiently correlate higher expression of *Aurkb* with the progression of human liver diseases.

### Upregulation of pT232-AURKB causes abscission failure in DEN-treated hepatocytes

Since activity and subcellular distribution play critical roles in modulating the functions of AURKB^45^, we next investigated whether DEN treatment affected the subcellular localization of AURKB. In hepatocyte primary cultures (Fig. 5a), pT232-AURKB showed typical subcellular distribution in control and DEN-treated hepatocytes during the cell cycle (Supplementary Fig. 5a). Consistent with previous findings, we observed that pT232-AURKB relocated from the central spindle to the midbody sequentially, pT232-AURKB levels gradually decreasing to be almost undetectable during abscission in control hepatocytes^43,44^. In stark contrast, DEN-treated hepatocytes showed higher levels of pT232-AURKB at all mitotic stages, compared to control hepatocytes (Supplementary Fig. 5a). Both AURKB and pT232-AURKB displayed higher intensity and wider distribution at the midbody in DEN-treated hepatocytes compared to control during abscission (Fig. 5b,c and Supplementary Fig. 5b,c). Additionally, longer intercellular bridges between daughter cells were observed in DEN-treated hepatocytes (Fig. 5b,d), resulting from the uncut intercellular bridge being pulled in different directions by the two daughter cells^13,47^. Regression analysis suggested that the length of intercellular bridge is positively correlated with the intensity of AURKB and pT232-AURKB at the midbody (Supplementary Fig. 5d). To test the causal relationship between the pT232-AURKB level at the midbody and intercellular bridges length, phosphorylation of AURKB was inhibited with AZD1152 (Barasertib), a highly specific AURKB inhibitor, in DEN-treated hepatocytes. A dose-dependent decrease of pT232-AURKB was confirmed in cultured hepatocytes under AZD1152 treatment (Supplementary Fig. 5e). Signal normalization of pT232-AURKB by AURKB suggested that only pT232-AURKB, not total AURKB levels, decreased under AZD1152 treatment at the midbody (Fig. 5b,c and Supplementary Fig. 5b), accompanied as hypothesized by a significantly reduced length of intercellular bridges in DEN-treated hepatocytes (Fig. 5b,d). Critically, the proportion of cultured hepatocytes failing abscission after DEN-treatment decreased significantly in the presence of AZD1152 (Fig. 5e). Altogether, these results show that DEN induces expression and activity of AURKB in mouse liver, facilitating abscission failure of hepatocytes.

**Figure 5.**
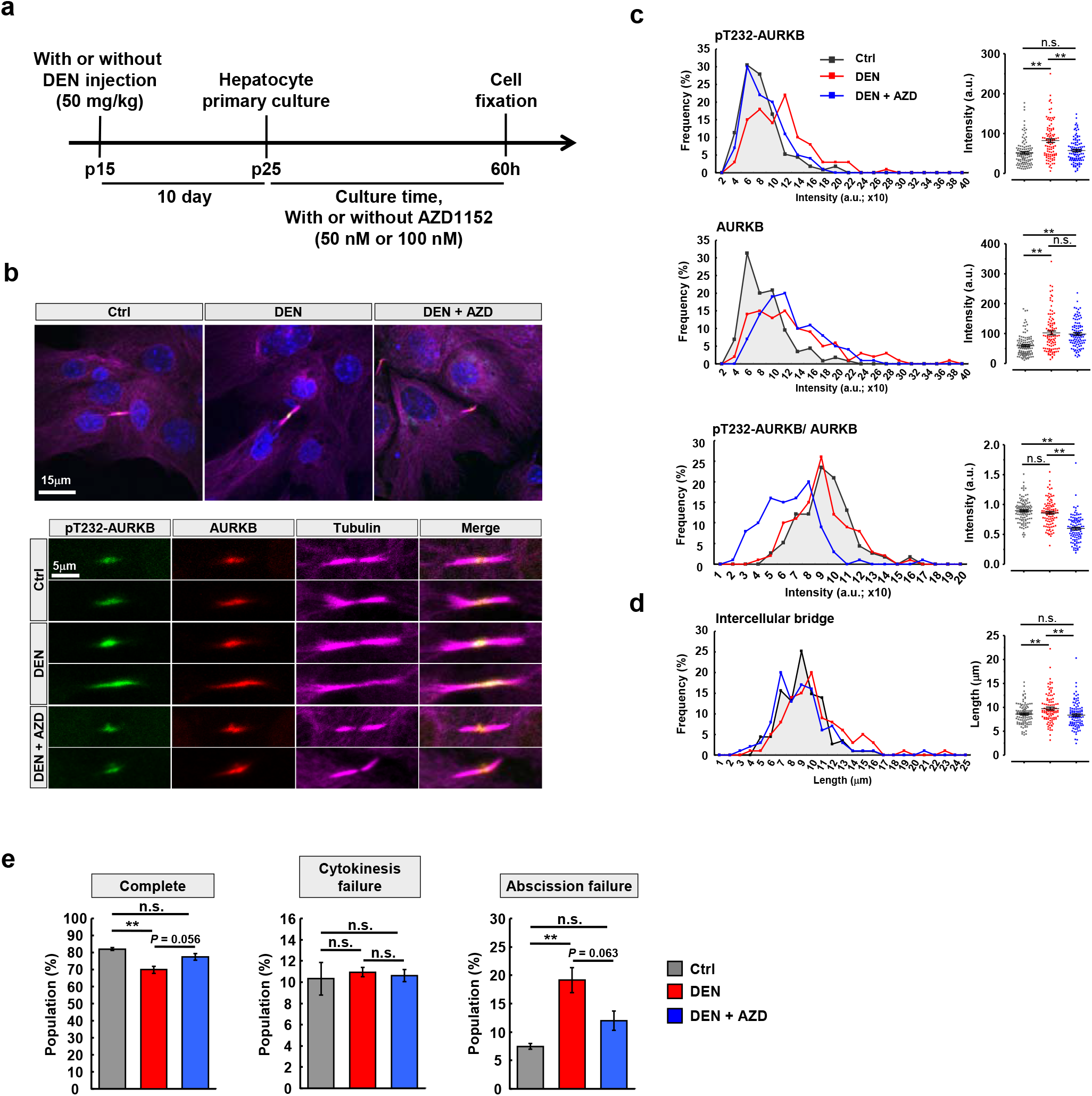
Hyperactivation of AURKB mediates DEN-induced abscission failure. (**a**) Schema for detection of AURKB and pT232-AURKB in hepatocyte primary culture: Injection of normal saline or DEN at p15 followed by mice sacrifice at p25 for primary culture. Cultured hepatocytes were treated with AZD1152 or equal amount DMSO. (**b**) Immunofluorescence of pT232-AURKB and AURKB during abscission in indicated cultured hepatocytes with or without 50 nM AZD1152 treatment. The morphology of nuclei and intercellular bridges were outline by counterstaining with tubulin and DAPI respectively. High magnification shows the detail structure of the midbodies. (**c**-**d**) Quantification of the intensity of pT232-AURKB, AURKB, and AURKB normalized pT232-AURKB at the midbody from Fig. 5**b**. The length of intercellular bridge was quantified by analyzed the highly condensed tubulin signal between daughter cells. n > 100 dividing hepatocytes at abscission stage per group from three independent experiments. (**e**) Quantification of the abscission failure ratio in the indicated cultured hepatocytes from time-lapse images. n > 500 dividing hepatocytes/group, three independent experiments/group. Statistic: One-way ANOVA, (**c**)-(**e**). Values represent the mean ± s.e.m., \**P* < 0.05, \*\**P* < 0.01, \*\*\**P* < 0.001. n.s., not significant.

### Reducing AURKB activity ameliorates DEN-induced hyperpolyploidy

High levels of pT232-AURKB have been reported to restrain abscission, leading to polyploidy through stabilization of intercellular canals^43,44^. We thus addressed whether hyperactivation of AURKB is involved in mediating DEN-induced nuclear enlargement *in vitro*. We first measured the size of nuclei during three consecutive days of primary hepatocyte cultures in the presence of vehicle, DEN or DEN + AXD1152 (Supplementary Fig. 6a), showing a progressive increase in all groups (Fig. 6a-c). The largest nuclei were observed in DEN-treated hepatocytes compared to control group, consistent with our previous data (Fig. 1c,d and 6a-c). Under DEN + AZD1152 treatment, nuclear size was reduced compared to DEN-treated hepatocytes, with AZD1152 showing a significant time-course effect (Fig. 6a-c). This result suggested that the level of pT232-AURKB plays a critical role in controlling hyperpolyploidization of hepatocytes.

**Figure 6.**
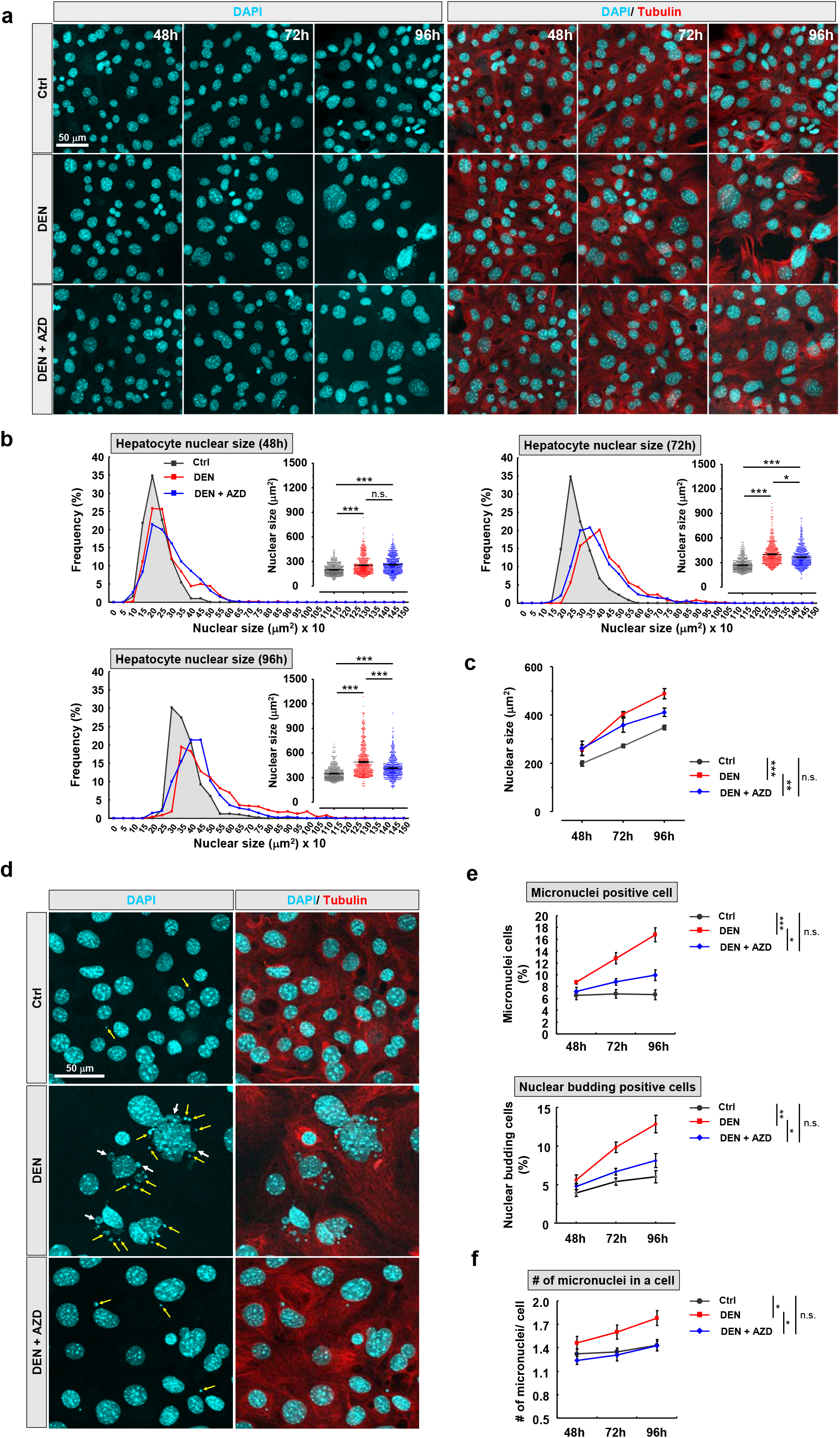
Hyperactivation of AURKB drives DEN-induced hyperpolyploidization of hepatocytes. (**a**) Pharmacological manipulations of AURKB activity, according to the schema in Supplementary Fig. 6a, changes the nucleus size of cultured hepatocytes. Cultured hepatocytes were treated with 50 nM AZD1152 or equal amount DMSO. Nucleus and cell morphology were outlined by DAPI (cyan) and tubulin (red) respectively. (**b**-**c**) Frequency distribution and dot plot graphs display the nuclear sizes of indicated cultured hepatocytes at different time points. Line graph shows the nucleus size in indicated cultured hepatocyte between 48h to 96h. n = 500 hepatocytes/group from four independent experiments. (**d**) Representative images show nuclear budding (white arrow) and micronuclei (yellow arrow) in indicated cultured hepatocytes. (**e**-**f**) The percentage of micronuclei and nuclear budding positive cell and the number of micronuclei in a cell of indicated cultured hepatocytes. Statistic: One-way ANOVA, (**b**); Two-way ANOVA, (**c**), (**e**), and (**f**). Values represent the mean ± s.e.m., \**P* < 0.05, \*\**P* < 0.01, \*\*\**P* < 0.001. n.s., not significant.

Because genomic reducing events have been demonstrated as a predictive sign of stemness and genetic instability during tumorigenesis^25^, we then performed a systemic survey of nuclear morphology. DEN-treated hepatocytes displayed higher micronuclei and nuclear budding ratio than in control group, AZD1152 treatment decreasing this ratio (Fig. 6d,e and Supplementary Fig. 6b). The number and size of micronuclei in DEN-treated hepatocytes were also increased, and significantly reduced after AZD1152 treatment (Fig. 6d,f and Supplementary Fig. 6c,d). Importantly, micronuclei and nuclear budding were also observed in CL hepatocytes with large nuclei in DEN-injected liver (Supplementary Fig. 6e), but were very rare in control liver, consistent with our *in vitro* results (Fig. 6d-f).

Together with our findings presented in Fig. 3, these results paint a picture in which aberrant AURKB-induced hyperpolyploidy also plays a role in the transformation of CL hepatocytes into preneoplastic cells via subsequent reduction of nucleus size, the sign of a probable genome reduction event. We further examined whether progeny cells with smaller nucleus and expressing cancer stem cell (CSC) markers could be detected in cultured hepatocyte after DEN treatment. Very few cells co-expressing the CSC markers EpCAM and Vimentin were observed *in vitro* but were significantly more abundant in hepatocytes cultured from DEN-treated liver compared to control, and decreased under AZD1152 treatment (Supplementary Fig. 6f,g). Altogether, our observations demonstrate that aberrant activation of AURKB is a critical pathway leading to DEN-induced hepatocyte hyperpolyploidization, promoting nuclear budding and micronuclei progression.

### Pharmacological blockade of pT232-AURKB reduces preneoplastic lesion formation in DEN-injected liver

We wondered whether halting hepatocyte hyperpolyploidization could be accompanied by attenuated preneoplastic lesions formation in DEN-treated liver. AZD1152 was injected intraperitoneally every two days starting one week after DEN injection for pathological examination (Supplementary Fig. 7a). DEN injection dramatically increased AURKB, but subsequent AZD1152 treatment reduced pT232-AURKB and its downstream target to control levels (Supplementary Fig. 7b). We next addressed the effect of AZD1152 on the number of BrdU-positive cells. The numbers of BrdU-positive hepatocyte and non-hepatocytes were analyzed, revealing AZD1152 significantly reduced the number of BrdU-positive hepatocytes in a dose-dependent manner, compared to DEN-treated liver. However, no significant effects of AZD1152 were seen on the proliferation of non-hepatocytes (Fig. 7a,b). Next, we examined the effect of AZD1152 on nucleus and cell size along the CV-PV axis. Like our previous findings (Fig. 1c,d and Supplementary Fig. 1c-e), DEN treatment caused hyperpolyploidization of CL and ML hepatocytes predominantly. Injections of AZD1152 and DEN significantly reduced the level of hyperpolyploidization (Fig. 7c-e and Supplementary Fig. 7c,d). We further investigated whether AZD1152 rescued the extent of nuclear vacuolation in DEN-treated liver. A higher frequency of nuclear vacuolation was detected DEN-treated liver; after AZD1152 treatment, however, the number and size of nuclear vacuolation were significantly reduced, which indicated the suppressive effect of AZD1152 on the DEN-induced appearance of the precancerous phenotype (Fig. 7f). To evaluate the suppressive effect of AZD1152 on preneoplastic lesions formation, we investigated the pathological morphology of liver tissues globally and microscopically. After 3 months of DEN injections, no obvious changes in the general morphology of the liver were seen compared to control liver. Small tumor nodules were sometimes observed on the surface of DEN-treated livers (Supplementary Fig. 7e), but not with AZD1152 (Supplementary Fig. 7e). We then examined H&E-stained liver sections microscopically, revealing that AZD1152 treatment significantly decreased the number of preneoplastic foci in DEN-treated liver (Fig. 7g). Moreover, AZD1152 treatment also partially reduced the size of the larger tumor nodules observed on the surface of the liver 6 months after DEN treatment (Supplementary Fig. 7f,g).

**Figure 7.**
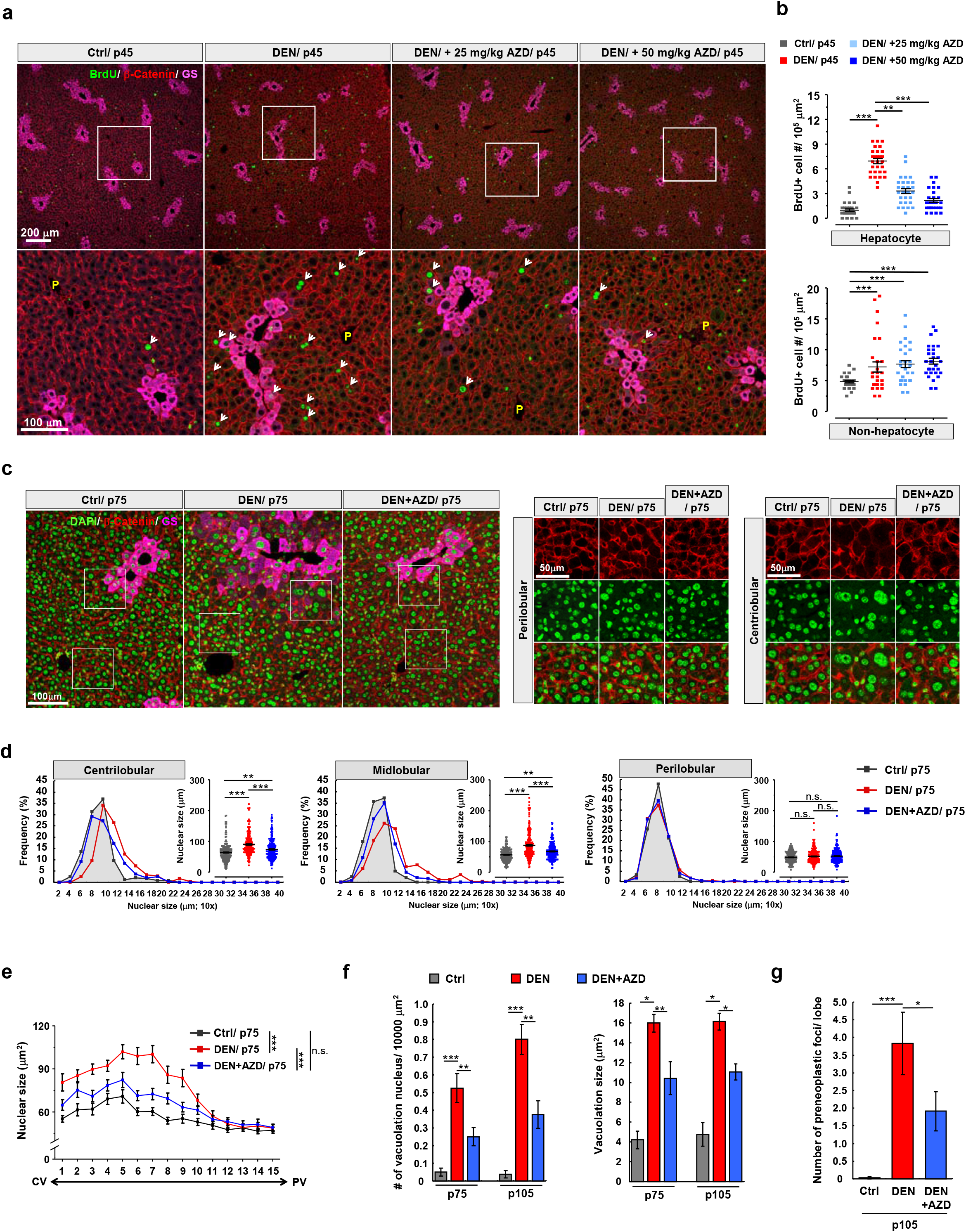
AZD1152 inhibits hyperpolyploidization of hepatocytes and preneoplastic foci formation in DEN-treated liver. (**a**) Immunohistochemistry of liver sections from indicated mice. The high magnification shows the detail information of BrdU positive hepatocytes. Schema of pharmacological manipulations was presented in Supplementary Fig. 7a. (**b**) The numbers of BrdU positive hepatocyte and non-hepatocytes in indicated liver tissues. N = 5 mice/group, and 6 different fields were selected randomly from each mouse. (**c**) Representative images from indicated liver tissues with 2 months of drugs treatment (DEN, 50 mg/kg; AZD1152, 25 mg/kg). The higher magnification shows the detail nucleus morphology nearby CL- and PL-regions. (**d**) The nucleus size of hepatocytes within specific liver lobule in indicated mice from Fig. 7C. N = 5 mice/group, and n = 750 hepatocytes/group. (**e**) Nucleus size of hepatocytes along CV-PV axis in indicated mice. N = 5 mice/group, and n = 750 hepatocytes/group. (**f**) The numbers of nucleus with vacuolation and vacuolation size in indicated liver after 2 and 3 months of drugs treatment (DEN, 50 mg/kg; AZD1152, 25 mg/kg). N = 5 mice for each group. (**g**) The numbers of preneoplastic foci in indicated liver with 3 months of drugs treatment (DEN, 50 mg/kg; AZD1152, 25 mg/kg). N = 5 mice/group. Statistic: One-way ANOVA, (**b)**, (**d**), (**f**), and (**g**); Two-way ANOVA, (**e**). Values represent the mean ± s.e.m., \**P* < 0.05, \*\**P* < 0.01, \*\*\**P* < 0.001. n.s., not significant.

Altogether, these data show that the DEN-induced emergence of preneoplastic hepatocytes at least partially depends on AURKB-mediated hyperpolyploidization. Critically, the transformation of hepatocytes into HCC cells can be prevented by AZD1152, once more underlining the potential of this inhibitor as a chemotherapeutic drug.

## DISCUSSION

Hepatocarcinogenesis is a multistep progression with genetic alterations that lead to malignant transformation of hepatocytes^3^, but how hepatocytes transformation occurs remains unresolved. Understanding the transformation process of hepatocytes into tumor cells is a critical issue to improve early clinical diagnosis of HCC and prevent hepatocarcinogenesis at an early stage. A significant feature observed in human and rodent liver tumor is a dramatic modification of the ploidy distribution of hepatocytes within the tumor nodules and in the noncancerous portions of the liver^21,31, 32,33^; however, these data were obtained at the cancerous stage, not in precancerous livers, offering little insight on the processes occurring during transformation. Here, we used a well-described HCC mouse model but focused on the initiation stage, demonstrating the mechanisms underlying hepatocyte transformation systemically (Fig. 8). Microscopic foci of preneoplastic lesions adjacent the CV marker were observed in DEN-injected liver, surrounded by hyperpolyploid hepatocytes (Fig. 1 and 2), in line with previous studies^28^. Deficiencies in circadian genes also increased abscission failure-mediated CL hepatocytes hyperpolyploidy and liver tumorigenesis^13, 48^, suggesting a possible link between the formation of hyperpolyploid hepatocytes and an early stage of HCC progression. Moreover, mouse models have established a relationship between stresses and pathological hyperpolyploidy in the liver^17,18,19,20^, but direct evidence of tumorigenesis are lacking. Our results reveal a molecular correlation between pathological hyperpolyploidy of CL hepatocytes and xenobiotics-induced HCC formation in mouse.

**Figure 8.**
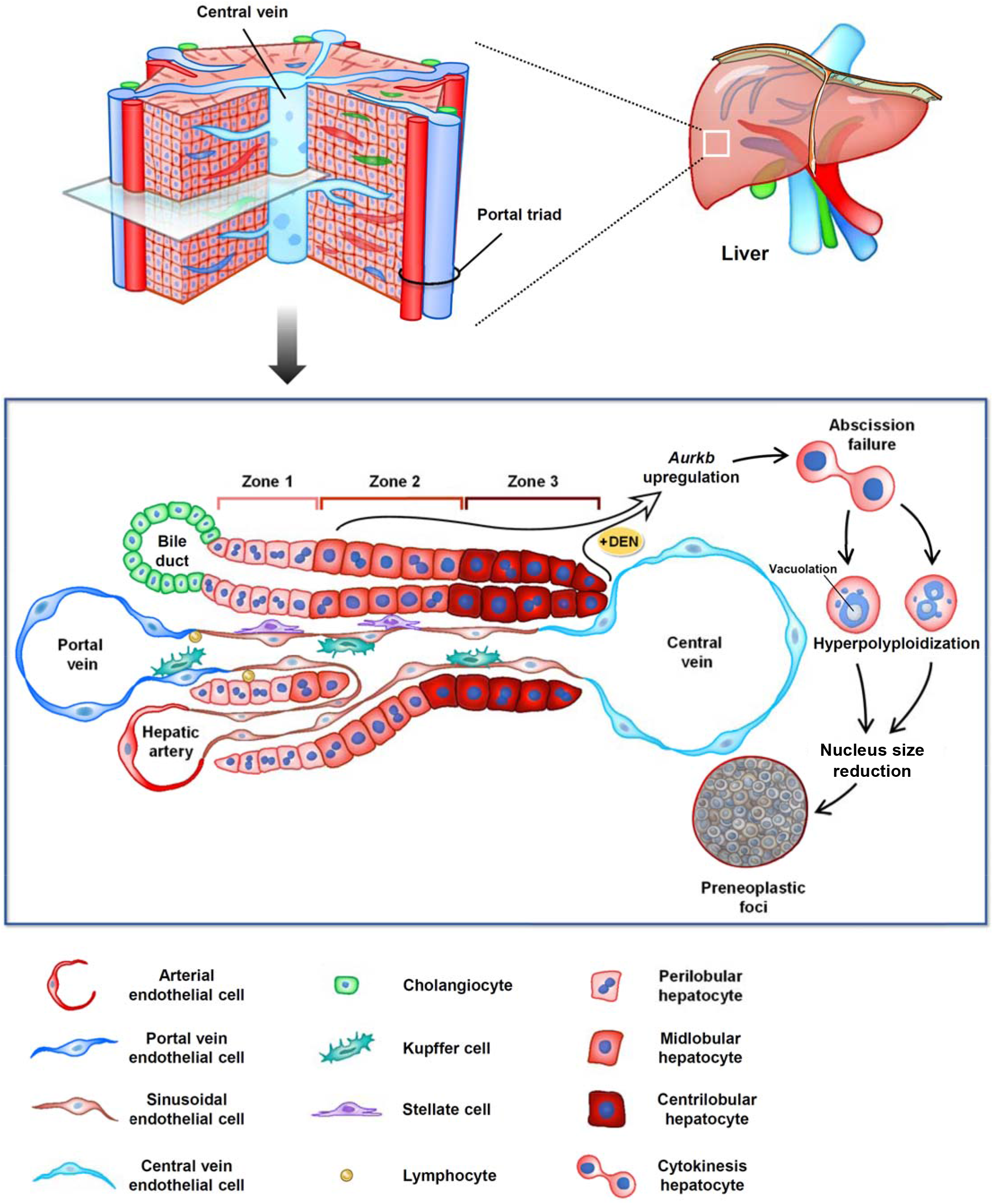
Schematic illustration of the generation of preneoplastic lesion from hyperpolyploid hepatocytes. Metabolically, hepatic lobule is composed by acinus that can be classified into three Zones: Zone 1 perilobule (PL), Zone 2 midlobule (ML), and Zone 3 centrilobule (CL). DEN treatment causes upregulation of *Aurkb* and abscission failure, which is a critical step for hyperpolyploidization of hepatocytes. The ratio of hyperpolyploid hepatocytes is increased predominantly in CL- and ML-region after DEN injection, accompanied with nuclear vacuolation and micronuclei. Hyperpolyploid hepatocytes might be the cell of origin in tumor initiation and subsequently transform into small size preneoplastic cells via an unknown process.

The expression and activity of AURKB are tightly controlled, either up- or down-regulation of AURKB causes chromosomal instability, abscission failure and polyploidization^45^. Upregulation of *Aurkb* is highly correlated with tumorigenesis of HCC^46,49,50^. Here, we have demonstrated that upregulation of AURKB plays a critical role in promoting abscission failure and hepatocyte hyperpolyploidization. Because endoreplication of hepatocytes has been also described under pathological conditions such as non-alcoholic fatty liver disease or liver cancer^12^, we cannot exclude the contribution of endoreplication in hepatocyte hyperpolyploidization.

According to *in vitro* and *in vivo* assays presented in Fig. 3g-l, our findings support the hypothesis that CL hepatocytes, after a reductive mitosis or other uncharacterized genome reduction events, may be the origin of HCC cells ^25,36,37^. A recent study indicated that “normal” polyploidy plays a tumor-suppressive role in the liver, but pathological hyperpolyploidy under stress conditions is a different situation^16^. We hypothesize that, under normal conditions, polyploidy indeed suppresses tumorigenesis. Under stress, hyperpolyploid hepatocytes may initially represent an adaptive and protective response. Beyond a bearable (and maybe reversible) level of stress and as genomic content further increase, hyperpolyploid hepatocytes may lose the ability to preserve their genomic stability, ultimately leading to a higher frequency of genomic alterations and tumorigenesis^24^. Indeed, the nucleus size of CL-enriched hyperpolyploid hepatocytes decreased, preceding the formation of tumor colonies (Fig. 3), echoing the findings of near-diploid HCC cells in human and rodent tumor nodules^31,32,33^. The proposed role of polyploid giant cancer cells in metastasis^25^ and the reversion of polyploid hepatocytes via the ploidy-conveyor^36,37^ also support the hypothesis of a genome reduction event in hyperpolyploid hepatocytes in the formation of preneoplastic lesions. Our results do not exclude the possible contribution of different stem/progenitor populations that may respond to oncogenic or growth-promoting signals secreted from hyperpolyploid hepatocyte. Although we did not formally investigate the mechanisms underlying genomic reduction, we observed multipolar dividing in CL hepatocytes-derived tumor cells (Fig. 3). Moreover, frequent nuclear budding and micronuclei in hyperpolyploid hepatocytes were identified *in vitro* (Fig. 6), and under DEN-treatment CSC markers-positive cells were often small and had small nuclei, a pathological phenotype that improved with AZD1152 (Supplementary Fig. 6e-g). Thus, based on our data, it is reasonable to speculate that hyperpolyploid hepatocytes, through reduction of genomic content, may play a source of stemness and tumor heterogeneity during initiation stage. The mechanisms underlying the transformation of hyperpolyploid hepatocytes into preneoplastic cells with low DNA content, and the impact of subgenomic content transmission remain to be investigated.

In conclusion, here we revealed that, after DEN-induced carcinogenic liver injury, hyperpolyploid hepatocytes arising from abscission failure are oncogenerative cells and a major source of preneoplastic lesions. These discoveries provide a significant contribution in the understanding of liver cancer biology, and may lead to new avenues in identifying potential pathological markers for early diagnosis and prevention of hepatocarcinogenesis.

## METHODS

For detailed materials and methods used in this study, please refer to the online supplementary materials and methods.

## ACKNOWLEDGEMENTS

The authors are indebted to Dr. Yi-Hsuan Huang and Dr. Ruey-Bing Yang for supplying facilities and drugs for *in vivo* and *in vitro* studies. The authors thank the pathology core of Institute of Biomedical Sciences (IBMS) for paraffin blocks and H&E stain processing. The authors also thank the assistance from Ms. Tzu-Wen Tai and Ms. Chia-Chen Tai in the flow cytometry core facility of IBMS (AS-CFII108-113), Academia Sinica for analysis of flow cytometry. The authors are grateful to Mr. Kuan-Yu Chou and Ms. Show-Rong Ma in microscopes facility of IBMS for images acquisition.

## AUTHOR CONTRIBUTIONS

Experimental conception and design: H.L., Y.-S.H., J.-M.F., M.D., H.C., and H.-W.C.; Development of methodology: H.L., Y.-S.H., J.-M.F., M.D., H.C., H.-S.H., and H.-W.C.; Acquisition of data: Y.-S.H., H.H. Lai., S.-H.L., Y.-L.L., P.-C.K., H.-S.H., H.-W.C., and P.-Y.Y.; Analysis and interpretation of data: H.L., Y.-S.H., J.-M.F., M.D., H.C., S.-H.L., and Y.-L.L.; Drafting of manuscript: H.L., Y.-S.H., and H.-W.C.; Critical revision: H.L., Y.-S.H., J.-M.F., M.D., H.C., and H.-W.C.; Study supervision: H.-W.C.; Equal contribution: These authors contributed equally to this article: ^10^, share co-first authorship; ^11^, share co-second authorship.

## COMPETING INTERESTS STATE

The authors declare no competing for financial interests.

## ETHICS APPROVAL

Animal studies were approved by the animal experimentation committee of Taipei Medical University and performed in accordance with the guidelines of the institutional committee for the use of animals for research. IACUC at TMU Approval No: LAC-20l6-0323.

