## Supplementary material & Figures for "Hyperpolyploidization of hepatocyte initiates preneoplastic lesion formation in the liver"

### Title

#### Funding

This work was supported by grants from the Taipei Medical University, Ministry of Science and Technology of Taiwan [MoST108-2628-B-038-002 and MoST106-2320-B-038-026].

#### Disclosure of Potential Conflicts Disclosure of Potential Conflicts

No potential conflicts of interest were disclosed.

#### Supplementary Material

##### Antibodies and chemicals

Antibodies used in detecting specific proteins are: BrdU (GeneTex, GTX26326), GAPDH (Santa Cruz, sc-25778), β-actin (Sigma-Aldrich, A5441), α-tubulin (Sigma-Aldrich, T5168), β-Catenin (Santa Cruz, sc-7199), Glutamine Synthetase (BD, 610518), AURKB (Abcam, ab2254), pT232-AURKB (Rockland, 660-401-667; Cell signaling, 2914), Histone H3 (ABclonal, A2348), pS10-Histone H3 (Cell Signaling, 9701), 9713) α-fetoprotein (Santa Cruz, sc-130302), Ep-CAM (Santa Cruz, sc-53532), Lamin B1 (ABclonal, A1910), γH2AX (ABclonal, AP0687), and mcherry (Abcam, ab167453). For immunofluorescence, AlexaFluor 488, 594 or 647-conjugated secondary antibodies are from Invitrogen. For immunoblot assay, horseradish peroxidase (HRP)-conjugated secondary antibodies were from Invitrogen. Chemicals used in the study are: Corning Matrigel Matrix (CORNING, 354234), ProLong antifade and DAPI reagents from Invitrogen. AZD-1152-HQPA (SML0268), Avertin (T48402), and diethylnitrosamine (55-18-5) were purchased from Sigma-Aldrich. BrdU (REF 000103/ LOT 1923353A; 3 mg/ml) was from Life technologies.

##### Experimental animals and drugs treatment.

All mice used for experiments were standard ICR strain exception of special cases. Female NSG™ immunodeficient mice (The Jackson Laboratory, stock No: 005557) with 8 weeks age were conducted for subcutaneous implantation. For high-fat diet treatment, wild type *C57BL/6J* mice were utilized, which are susceptible to diet-induced obesity. Mice were maintained in 12h light/ 12h dark cycle (LD) with food and water *ad libitum*. Before perfusion and sacrifice, mice were anesthetized with Avertin (0.25 mg/g body weight). For observation of DEN-induced preneoplastic foci formation, mice were double injected (i.p.) with DEN (50 mg/kg body weight) at 15 days and 22 days age and sacrificed at indicated times. For Aflatoxin B1 (AFTB1) model, male mice were double injected with a 6 mg/kg body weight dose of AFTB1 dissolved in DMSO at 15 days and 22 days age and sacrificed at indicated times. For $CCl_4$ chronic-injury model, male mice with 15 days age were injected with a 0.5 mg/kg body weight dose of $CCl_4$ dissolved in corn oil every 3 days for 10 doses to induce chronic damage. For high-fat diet model, control and experimental groups were treated with 8 kcal% normal chow diet and 45 kcal% high-fat diet at 8 weeks age for indicated times, respectively. For time-lapse recording of hepatocytes *in vitro*, single dose injection of DEN (50 mg/kg body weight) was applied through i.p. to 15 days old mice, and DEN-injected mice were sacrifice at 25 days age for hepatocyte primary culture. To suppress AURKB activity *in vivo*, AZD1152 (25 or 50 mg/kg body weight) was injected one day after DEN injection through i.p. every two days until sacrifice. AZD1152 powder was dissolved in DMSO to get the stock with concentration of 10 mg/ ml. Then, AZD1152 was finally diluted into working concentration by normal saline for i.p. injection. For cell proliferation analysis *in vivo*, BrdU (10 μl/g body weight; 3 mg/ml) purchased from company was directly used and injected i.p. for 5h. All animal experiments were approved by the animal experimentation committee of Taipei Medical University and performed in accordance with the guidelines of the institutional committee for the use of animals for research. The different groups were housed together in the same cages in all animal experiments.

**Human sample analysis**

To make a link between biomedical discovery in mouse model and clinical findings, genomic alterations of *Aurkb* in various clinical cases were obtained from Gene Expression Omnibus (GEO) datasets in NCBI. Genomic expression profiles of human liver biopsies were extracted from Series Matrix Files of GEO datasets (http://www.ncbi.nlm.nih.gov/geo/). The fold change of *Aurkb* in patients with steatosis, NAFLD, and various stage of fibrosis was acquired from GSE89632 and GSE66232. Examination of *Aurkb* expression in tumor and non-tumor biopsies of human liver was obtained from GSE20140, GSE54236, and GSE64041. The correlation between *Aurkb* expression level and survival time of patients was analyzed according to GSE54236. The case numbers and statistical analysis were display in the figure legends.

**Hepatocyte Primary Culture**.

Primary mouse hepatocytes were isolated from 3-week-old mouse livers using a two-step collagenase I perfusion protocol. Briefly, liver was perfused with $Ca^{2+}$ and $Mg^{2+}$ free Hank's balanced salt solution (HBSS) from inferior vena cava with a flow rate of 5 ml per minute for 3 min at RT. An incision was made in the portal vein to let blood out of the liver. Liver was then perfused with 10 mg/ml collagenase I (Worthington Biochemical Corporation) in $Ca^{2+}$ and $Mg^{2+}$ positive HBSS (Gibco) with a flow rate of 3 ml per minute for 10 min. After perfusion, liver was disrupted with scissors and dissociated by pipetting gently in DMEM medium. Cell suspension was filtered through a cell strainer (100 $\mu m^2$) and mixed with equal volume of 90% Percoll (Sigma) in HBSS. To collect live hepatocytes, the mixture was centrifuged at 100× *g* for 5 min, and the pellet was resuspended in 199 medium (Gibco) containing 5% FBS. Isolated hepatocytes were seeded on collagen-coated 6-well plate (density = $2 \times 10^5$) or 12-well plate containing collagen-coated coverslips (density = $5 \times 10^4$) and incubated at 37°C with 5% $CO_2$. Medium was refreshed after 4h of seeding. After 24h of seeding, medium was replaced by 199 medium containing 10 mM HEPES (Gibco), 4.5 mg/ml glucose, 2 mM L-glutamine (Invitrogen), antibiotic-antimycotic (Gibco), 20 mM sodium-pyruvate (Gibco), 5 ng/ml sodium selenite (Sigma), 5 mM nicotinamide, 10 mg/ml transferrin (Sigma), 10 mM 3,3',5-triiodo-L-thyronine sodium salt (Sigma), 50 ng/ml recombinant mouse EGF (Invitrogen), 100 nM insulin (Sigma), 100 nM dexamethasone (Sigma), and 5% FBS. Medium was refreshed every 24 h throughout the culture time.

**Time-lapse Recording**.

The cytokinetic structures of hepatocytes were monitored *in vitro* by time-lapse microscopy (Leica, DMI 6000B) equipped with auto-focus system (MAC 6000 system) and live cell instrument (CU-109, $CO_2$/ air gas mixer FC-5). Primary cultured hepatocytes isolated from the livers of 3-week-old mice were used since the proliferation ability of hepatocytes at this age is most vigorous and over 90% of hepatocytes are diploid and tetraploid[13]. 24h after seeding, images of hepatocytes were recorded and cellular behavior and structures were visualized by phase contrast system with HC PL Fluotar 20x/0.50 PH2 dry objective lens and 100 msec exposure time. Hepatocytes were incubated in a chamber maintained at 37°C and

5% $CO_2$. Images were taken at 10 min intervals for approximately 150h by Andor iXon EMCCD Camera. The optimal focal plane was set at beginning of each image session and adjusted by auto-focus system with 5-second intervals throughout the image recording. The recorded images were further analyzed via ImageJ software.

**Isolation of centrilobular and perilobular hepatocytes**

Enrichment of hepatocytes from CL or PL region was performed by using the digitonin-collagenase perfusion system as previously described[31-32]. Mice were anesthetized with Avertin (0.25 mg/g body weight) before liver perfusion. For isolation of centrilobular hepatocytes, 24G i.v. catheters (Terumo) were set into portal vein and inferior vena cava, and blood was removed by perfusion with $Ca^{2+}$ and $Mg^{2+}$ free HBSS from portal vein with a flow rate of 5 ml per minute for 3 min at RT. Digitonin buffer (4 mg/ml in $Ca^{2+}$ and $Mg^{2+}$ free HBSS) was infused at a rate of 10ml/min at RT from portal vein until a regularly scattered periportal discoloration pattern emerging on the surface of liver (this took 10 to 30 seconds). Digitonin was then removed by retrograde perfusion with $Ca^{2+}$ and $Mg^{2+}$ free HBSS containing 1mM EGTA from inferior vena cava for 2 min with a flow rate of 5 ml/min. Retrograde infusion of 1 mg/ml collagenase I in $Ca^{2+}$ and $Mg^{2+}$ positive HBSS from inferior vena cava with a flow rate of 3 ml per minute for 10 min was performed subsequently. After 10 min of collagenase perfusion, moved digested liver into DMEM medium, and liver was disrupted with scissors followed by dissociation of hepatocytes with pipetting. Cell suspension was filtered through a cell strainer (100 $\mu m^2$) and mixed with equal volume of 90% Percoll (Sigma) in HBSS to collect live centrilobular hepatocytes. For isolation of perilobular hepatocytes, blood was removed by perfusion with $Ca^{2+}$ and $Mg^{2+}$ free HBSS from central vein followed by digitonin buffer infusion. Digitonin was then removed by retrograde perfusion with $Ca^{2+}$ and $Mg^{2+}$ free HBSS containing 1mM EGTA from portal vein, which is followed by infusion of 1 mg/ml collagenase I in $Ca^{2+}$ and $Mg^{2+}$ positive HBSS from portal vein for 10 min. Subsequently, live perilobular hepatocytes were isolated by Percoll-dependent centrifugation method for further inoculation.

**Tumorsphere assay and colony formation assay**

Centrilobular and perilobular hepatocytes were isolated by the digitonin-collagenase perfusion system from control and one month DEN-treated mice. Cells were resuspended in 199 culture medium containing 10 mM HEPES, 4.5 mg/ml glucose, 2 mM L-glutamine, antibiotic-antimycotic, 20 mM sodium-pyruvate, 5 ng/ml sodium selenite, 5 mM nicotinamide, 10 mg/ml transferrin, 100 nM insulin, 100 nM dexamethasone, 10 mM 3,3',5-triiodo-L-thyronine sodium salt, 50 ng/ml recombinant mouse EGF, 20 ng/ml bFGF, 2% B27 supplements (Gibco), and 1% N2 supplement (Gibco). Cells were then cultured in Ultra-low non-attachment 24-wells plate (Corning, 3473) with a serial dilution including 500, 1000, and 2000 cell number at 37 °C and with 5% $CO_2$. The medium was refreshed every week to prevent disturbance the formation of the tumorspheres. After one-week incubation, pipetting was performed gently with 1 ml tips to dissociate aggregated cells, and tumorspheres were monitored by phase-contrast microscope (Leica, DMI 6000B) with HC PL Fluotar 20x/0.50 PH2 dry objective lens. Numbers of tumorsphere (diameter > 100 μm) were analyze by ImageJ. For colony formation assay, Ultra-low non-attachment plate was replaced by regular 6-well plate (Greiner CELLSTAR, 657160) with Type I collagen coating. After 21 days inoculation, cells were washed with PBS and then fixed by 4% PFA in PBS for 1 h at RT, which is followed by crystal violet staining (0.5% w/v) and counted using a stereomicroscope.

**Subcutaneous implantation**

Hepatocyte spheres derived from CL or PL hepatocytes were conducted for subcutaneous injection. Spheres derived from 2000 numbers of hepatocytes were used for implantation. During the subcutaneous implantation, spheres in 100 μl 50% Matrigel/media without serum was injected into subcutaneous region of skin. Spheres derived from PL- and CL-enriched hepatocytes were injected into the left and right flank under the skin of the dorsal body surface of each mouse respectively. Mice were monitored daily in the first 10 days of post-injection and when the tumor started to grow, the tumor size was measured every five days. Tumor size was measured using a caliper. When the tumor size reached around 1 $cm^3$, mice were euthanized and the tumor was harvested.

Definition of preneoplastic lesion

The sequentially pathological changes have been well described in human and rodents, and the terminology and criteria for various lesions has been defined based on consensus diagnosis. The first visible pathological lesion in the progression of cancer can be classified as a preneoplastic lesion, characterized by as a small focus of cellular alternation and appear to represent clonal expansion of about a hundred cells. These preneoplastic lesions are difficult to detect with the naked eyes but can be identified microscopically. For tumor nodules, constituted by continuously growing of preneoplastic cells, can be observed directly through naked eyes on the tissue surface. Tumor nodule may identify as benign tumor since they grow by expansion of preneoplastic cells and are not invasive with well differentiated morphology. For instance, in Figure 1b, the whitish and round clump on the liver surface can be defined as a tumor nodule.

**Immunohistochemistry and H&E staining**

After anesthetization, mice were perfused with cold PBS followed by 4% paraformaldehyde (PFA)/ PBS. Specimens were post-fixed with 4% PFA/ PBS and embedded in paraffin-wax with standard protocol. Five-micrometer-thick Paraffin sections were prepared as 10 μm thickness and deparaffinized with xylene. Rehydration was performed with a serial concentration ethanol followed by antigen retrieval in Tris–EDTA buffer (pH 9.0) for 5 min with pressure cooker. Sections were immersed in PBS containing 0.2% Triton X-100 for 30 min at RT for permeablization. For immunofluorescence labelling, sections were treated with Super Block and mouse to mouse block (ScyTek Lab) for 10 min and 60 min followed by blocking with 5% BSA and 5% FBS in PBS for 1h. After blocking, sections were treated with primary antibodies diluted in PBS containing 1% BSA, 1% FBS, and 0.05% Triton X-100. Following overnight incubations at 4°C, sections were washed extensively with PBS containing 0.1% Triton X-100 and then incubated with secondary antibodies conjugated with Alexa fluorophores and DAPI. After PBS wash, sections were mounted in Prolong Gold mounting medium with anti-fade reagent (Invitrogen) for image acquisition. Histological staining with hematoxylin and eosin was performed by the institutional Pathology Core staff. Sections after the removal of paraffin and rehydration were stained with hematoxylin and eosin according H&E staining standard protocol.

**Immunocytochemistry**

Hepatocytes inoculated on collagen-coated coverslips were washed with PBS and then fixed with 4% PFA/ PBS at RT for 10 min. Permeablization was performed with 0.2% TX-100 in PBS at 4℃ for 20 min. Hepatocytes were then conducted for blocking with 5% BSA and 5% FBS in PBS for 1h at RT. For immunofluorescence labeling, hepatocytes were treated with primary antibodies diluted in PBS containing 1% BSA, 1% FBS, and 0.05% Triton X-100. Following overnight incubations at 4°C, hepatocytes were washed extensively with PBS containing 0.05% Triton X-100 and then incubated with secondary antibodies conjugated with Alexa fluorophores and DAPI. After PBS wash, cover slips were mounted in Prolong Gold mounting medium with anti-fade reagent (Invitrogen) for image acquisition.

**Image acquisition**

For evaluation of nuclear and cellular size in the liver, DAPI labeling and anti-β-catenin antibody staining were applied to outline nuclear and cellular architecture, respectively. Immunostained tissues were observed with a confocal laser scanning microscope LSM780 (CarlZeiss, Germany) equipped with an argon laser (excitation 488nm) and a DPS laser (excitation 561nm). Serial optical Z-sections were acquired using a Plan Apochromat

10X/0.45 M27 objective with 2048x2048 resolution. The x-y plane resolution as calculated by scaling 0.629 μm with apertures 29 μm. For observation of signals at midbody region, serial optical Z-sections were acquired using a Plan Apochromat 20X/0.8 M27 objective with 2048x2048 resolution. The x-y plane resolution as calculated by scaling 0.231 μm with apertures 27 μm. The nuclear budding of hepatocytes was detected by serial optical Z-sections using a Plan Apochromat 40X/1.4 Oil DIC M27 objective with 2048x2048 resolution. The x-y plane resolution as calculated by scaling 0.148 μm with apertures 29 μm. Projection images were generated using ZEN black image analysis software (Carl Zeiss, 2011 SP7 FP3, version 14.0.0.0) with full resolution and shown for architecture of hepatocytes *in vivo* and *in vitro*.

**Images quantification**

For the nuclear and cellular size measurement alone CV-PV axis, the CV-PV axis was subdivided into 15 parts (Supplementary Fig. 1a), and the less number indicates the region closed to central vein region[13]. The traditional 3-zone classification of hepatic lobule divides CV-PV axis into three equal parts: namely, centrilobular zone corresponds to 1–5, midlobular zone to 6–10, and periportal zone to 11–15. Only hepatocytes (cells with round or oval nuclei) were analyzed in vivo and in vitro. Nuclear and cellular sizes were measured according to DAPI and β-catenin signal respectively. Data were obtained from three to five mice for each group. For quantification, at least five different fields of the microscope were selected randomly from each mouse liver, and approximately 50 CV-PV axises were analyzed for quantification. Mono- and bi-nucleated hepatocytes were distinguished by comparing DAPI and membrane-labeling images.

The length of intercellular bridge between daughter hepatocytes was quantified based on the β-tubulin signa[13]. Intercellular bridge is a region with highly condensed microtubules, and it is defined as a transient structure with about 1 to 2 μm in diameter only appearing towards the end of cytokinesis and just prior to the complete separation of nascent daughter cells.

Length of the brightly labeled tubulin bundle in the center of the intercellular bridge between nascent daughter cells was measured using ImageJ. The subcellular fluorescent intensity and length of AURKB and pT232-AURKB during abscission were quantified by background-corrected line scans along the central spindle region or midbody using ImageJ. The integrated fluorescence intensity along this line was used as the estimated amount of specific protein in this region. Over 100 dividing hepatocytes were analyzed for each group from three independent experiments. All images were acquired with a fixed exposure time and condition in the same experiment. All quantification of fluorescence intensities was performed under raw 16-bit images, and statistical analysis was performed by GraphPad Prism-5.0.

For quantification of the size and numbers of budding nuclei and micronuclei in cultured hepatocytes, DAPI and β-tubulin signals were conducted to outline nuclear and cellular morphology of hepatocytes respectively. Nuclear budding was defined as a protrusion part from regular nuclei, and micronuclei was characterized as a small separated part with DAPI positive signal. The size and numbers of budding nuclei and micronuclei were analyzed by ImageJ. Over 150 cultured hepatocytes with nuclear budding and micronuclei were analyzed for each group from at least three independent experiments.

Quantitative analysis of cell division types from time-lapse images of hepatocytes in culture was performed by ImageJ. The complete cytokinesis, cytokinesis failure, and abscission failure are defined as previous described[13]. Complete cytokinesis is defined as two individual daughter cells generated from single hepatocyte after bipolar dividing. For cytokinesis failure, hepatocytes perform nuclear division (karyokinesis) only, and no contractile ring formation and cytokinesis are progressed. For abscission failure, ingressed cleavage furrow emerges between two daughter cells; however, the daughter cells lose its ability to cut off intercellular bridge, and furrow regression is performed to generate a single cell with multiple nuclei.

**Flow Cytometry**.

Hepatocytes were isolated by collagenase perfusion system and resuspended in ice-cold PBS at a density of 2 x $10^6$ cells/ ml. Cells were fixed with 4% PFA at 4°C overnight with gentle rotation. Cells were washed with PBS containing 0.25 mg/ml RNase A and stained with 2 μg/ml DAPI at 4°C for 60 min. DNA content of labeled cells was measured by flow cytometer (Becton Dickinson, LSRII SORP – 17 color analyzer). Acquired data were analyzed by BD FACSDiva software v6.2

**Quantitative Real-Time PCR**

Total RNA in liver was purified by TOOLSmart RNA Extractor (BioTools, DPT-BD24) and RNeasy Mini kit (QIAGEN, 74104). The isolated RNAs were reverse transcribed by using an oligo-dT/random primer mixture and ImPromII Reverse Transcriptase (Promega) with equal amount of purified RNA. Quantitative(q) PCR involved the Universal Probe Library and Lightcycler 480 system (Roche). The comparative Ct (threshold cycle value) method was used to calculate relative expression. Raw data were normalized by the expression level of *Actb*, which showed no significant difference in both control and drug treated mice. The primer pairs used in qPCR are list in Table 1.

**Immunoblot assay**

Protein extracts were prepared from equal amounts of the liver or cultured hepatocytes in Laemmli sample buffer supplemented with 1X protease inhibitor cocktail (Roche) and phosphatase inhibitors (Tocris). Homogenates were sonicated by ultrasonic cell disruptor (Misonix Sonicator 3000) with power level 2 to disrupt genomic DNA, and insoluble debris were removed by centrifugation at 14,000 x*g* and 4℃ for 10 min. Protein lysates were boiled at 55°C for 15 min and then resolved by 10% SDS-PAGE mini-gels, transferred to nitrocellulose membranes. The membranes were then blocked in 5% skim milk in 1X TBST (TBS with 0.1% Tween 20) for 60 min, followed by the incubation of indicated primary antibodies at 4℃ overnight. Specific horseradish peroxidase-conjugated secondary antibodies were used for enhanced chemiluminescence detection (ECL-Prime, GE Healthcare Life Sciences) by Image Quant LAS 4000 (Fujifilm).

**Statistical analysis**

Mice and cultured hepatocytes were randomly assigned for time-course study and drugs treatment. Imaging fields were randomly selected during image acquisition. The sample size among experimental groups was kept as equally as possible. The experiments and analysis were conducted in a blind manner and replicated at least three times independently. For the *in vivo* studies, at least five mice were used for each group. GraphPad Prism 5.0 software was applied to produce the graphs and statistical analysis.

We conducted experiments including BrdU positive nuclear area (Supplementary Fig. 1j), nuclear size between vacuolation and non-vacuolation hepatocytes (Fig. 2b), nuclear to cytoplasmic ratio (Fig. 2h), nuclear and cytoplasmic size between tumor and non-tumor cells (Fig. 3a), nuclear size of colony and non-colony cells (Fig 3i), nuclear size of region specific hepatocytes (Supplementary Fig. 3b), tumorsphere size (Supplementary Fig. 3f), *Aurkb* gene expression in HCC patients (Fig. 4f) to two-tailed unpaired Student's t-test with Welch correction. Supplementary Fig. 4c was analyzed statistically by two-tailed paired Student's t-test. For the ratio of different cell division type (Fig. 1i), qPCR analysis of CL and PL markers expression (Fig. 3e), and expression of cytokinesis genes (Fig. 4c), two-tailed Mann Whitney nonparametric test was applied.

For the effect of time on nuclear, cellular and micronuclear size after drugs treatment (Fig 1d, Fig. 6b, Fig. 7d, Supplementary Fig. 1f, dot plot graphs of Supplementary Fig. 6c and 6d, and Supplementary Fig. 7c), numbers of BrdU positive hepatocytes (Fig. 1f), distribution of vacuolation nuclei and tumor foci (Fig. 2d and i), colony numbers (Fig 3g), *Aurkb* gene expression in various liver disease patients (Fig. 4e), the intensity of pT232-AURKB and AURKB signals at the midbody (Fig. 5c), the length of intercellular bridge, pT232-AURKB and AURKB signals (Fig. 5d, Supplementary Fig. 5c), the ratio of different cell division type after drugs treatment (Fig. 5e), the change of nuclear size after drugs treatment (Fig. 6b), the effect of AZD1152 on the numbers of BrdU positive cells, nuclear size of hepatocytes, numbers and size of vacuolation in hepatocytes, and preneoplastic foci numbers (Fig. 7b, d, f and g), the numbers of EpCAM and Vimentin double positive cells (Supplementary Fig. 6g), and the tumor diameter and tumor numbers in Fig. 7f were statistically analyzed by One-way ANOVA.

Two-way ANOVA was applied to analyze the distribution of BrdU positive cell along CV-PV axis at different stages of DEN treatment (Fig. 1g and Supplementary Fig. 1i), the effect of DEN on the numbers of vacuolation and preneoplastic foci at different time-courses (Fig. 2c and e), *in vivo* tumor formation rate (Fig 3j), the effect of region specific hepatocytes on tumorsphere formation (Supplementary Fig. 3e), the effect of AZD1152 on the size of nuclei and micronuclei, percentage of cells with micronuclei and nuclear budding, and the numbers of micronuclei in a cell at different time points (Fig. 6c, e, and f, and the line graphs of Supplementary Fig. 6c and d), and the effect of drugs on the frequency distribution of nucleus and cell size along CV-PV axis (Fig. 7e, and Supplementary Fig. 7d). Bonferroni's multiple comparison test was applied for comparisons among multiple conditions following One-way or Two-way ANOVA tests.

Correlation quantifies the degree to which two variables are related including expression level of *Aurkb* and the survival time and tumor doubling time of patients (Fig. 4h and Supplementary Fig. 4d), nuclear size and the numbers and size of vacuolation (Supplementary Fig. 2c), the length of intercellular bridge and AURKB intensity (Supplementary Fig. 5d) were statistically analyzed by correlation coefficient. Kaplan Meier Survival Analysis was conducted to statistically analyze the overall survival curves of patients with different *Aurkb* expression (Fig. 4g). Sample numbers and statistical results were indicated in the figure legends precisely. Data were presented as the mean ± standard error (s.e.m.), and P values less than 0.05 were considered significant. P values are represented as *$P < 0.05$, **$P < 0.01$ and ***$P < 0.001$. n.s., not significant; n.d., not detectable.

### Supplementary Figures

#### Supplementary Figure 1

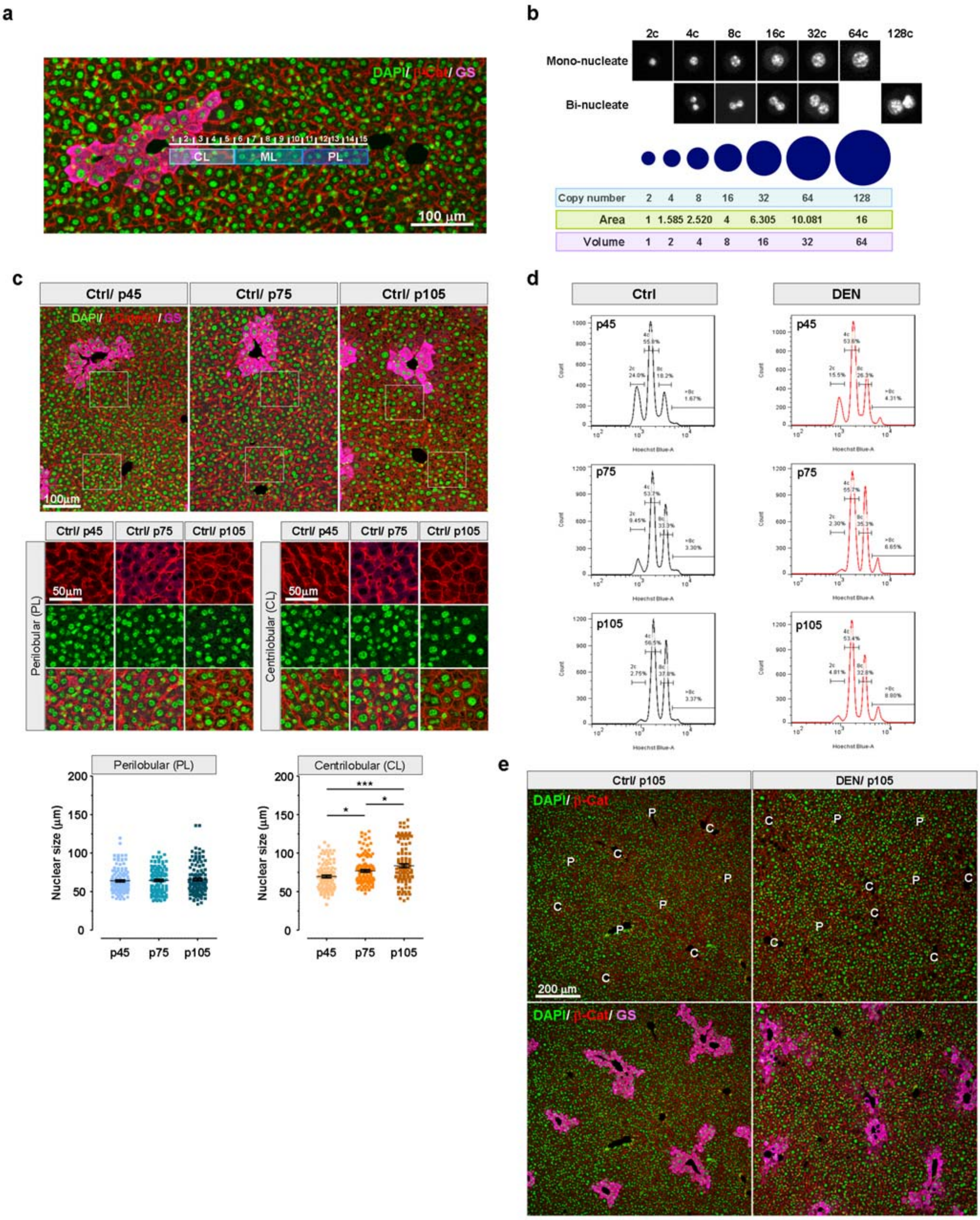

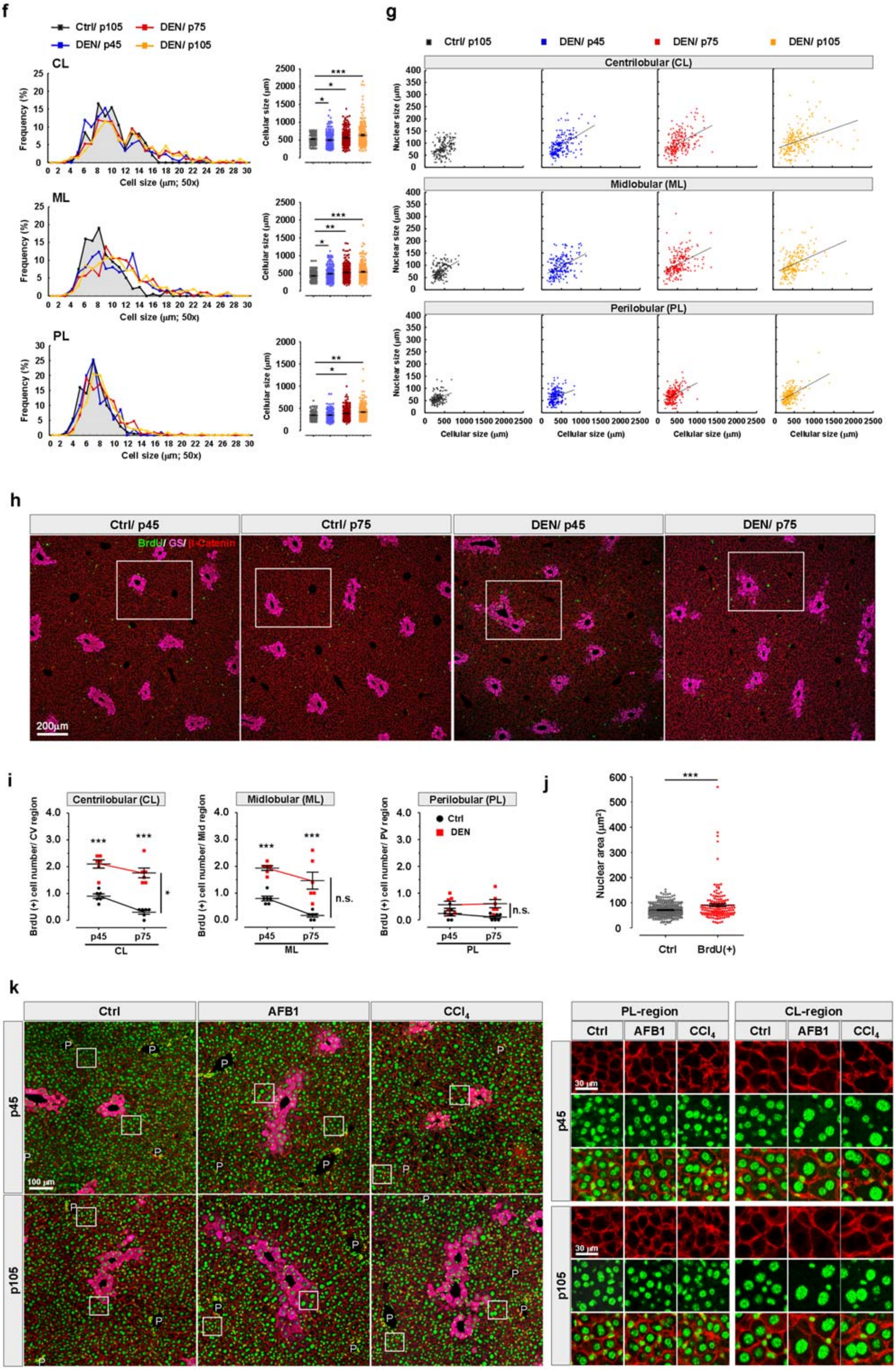
f
Ctrl/ p105
DEN/ p75
DEN/ p45
DEN/ p105
CL
ML
PL
Frequency (%)
Cell size (μm; 50x)
Cellular size (μm)
g
Centrilobular (CL)
Midlobular (ML)
Perilobular (PL)
Nuclear size (μm)
Cellular size (μm)
h
Ctrl/ p45
Ctrl/ p75
DEN/ p45
DEN/ p75
BrdU/ GS/ β-Catenin
200μm
i
Centrilobular (CL)
Midlobular (ML)
Perilobular (PL)
BrdU (+) cell number/ CV region
BrdU (+) cell number/ Mid region
BrdU (+) cell number/ PV region
Ctrl
DEN
n.s.
j
Nuclear area (μm²)
Ctrl
BrdU(+)
k
Ctrl
AFB1
CCl4
p45
p105
100 μm
30 μm
PL-region
CL-region

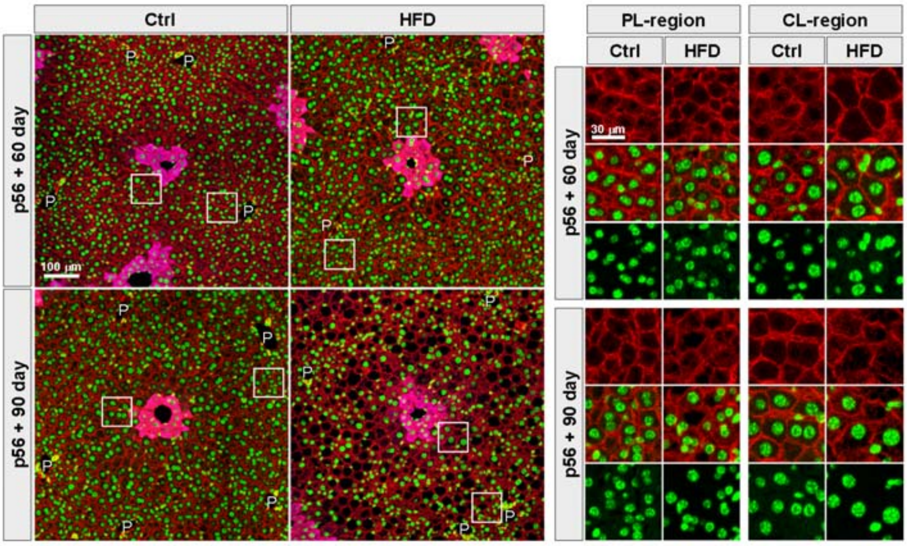

**Supplementary Figure 1.** DEN treatment causes pathological hyperpolyploidy of hepatocytes within CL and ML regions. (**a**) Example of image displays subdividing of CV-PV axis into 15 parts for further analysis. Classic hexagonal shaped liver lobule with three specific zones including centrilobular (CL), midlobular (ML) and perilobular (PL) regions, and each specific zone is composed by 5 parts as indicated. Liver slice was co-stained with DAPI (green) and β-catenin (red), and CL regions marker GS (magenta). (**b**) Representative images show the nuclear morphology of hepatocytes isolated from 3 month DEN-treated mouse liver and counterstained with DAPI. Assuming that the concentration and distribution of nuclear chromatin are equal and similar in every hepatocyte, and it follows Lambert-Beer's law during image acquisition, the relationship between the cross-sectional area of the spherical nucleus and the ploidy can be shown as below. (**c**) Representative immunostaining of liver sections from control mice at indicated ages. High power images show the cell and nuclei size of hepatocytes within CL and PL region. Dot plot graphs illustrate the changes of cell and nuclei size of hepatocytes during liver development in control mice. Mouse number (N) = 3 per group, and cell number (n) = 120 per group. (**d**) Representative histograms of flow cytometry demonstrate the DAPI-labeled DNA content of control and DEN-treated hepatocytes at indicated ages. DEN-treated hepatocytes showed more populations with higher DNA content than control hepatocytes as comparing at the same age. (**e**) Images of liver sections from control and 3 month DEN-treated liver under low magnification. The enlarged nuclei were observed in DEN-treated liver nearby CL and ML region of liver specifically. Liver slices were co-stained with DAPI (green) and β-catenin (red), and CL regions marker GS (magenta). (**f**) The quantitative data show the cell size of hepatocytes along CV-PV axis displayed as frequency distribution and dot plot graphs. Mouse number (N) = 5 per group, and cell number (n) = 750 per group. (**g**) The scatter plot graph displays the relationship between nucleus and cell size in control and DEN-treated liver at indicated times. The bigger nucleus and cell size were detected in DEN-treated liver with time-dependent manner. Mouse number (N) = 5 per group, and cell number (n) = 750 per group. (**h**) Representative images show the distribution of BrdU in control and DEN-treated liver under lower magnification at indicated time. Boxed regions are shown in Fig. 1e with high magnification. Liver slices were co-stained with BrdU (green) and β-catenin (red), and CL regions marker GS (magenta). (**i**) Quantitative data display the numbers of BrdU positive hepatocytes at two developmental time points in control and DEN-treated livers. Higher numbers of BrdU positive hepatocytes were detected in DEN-treated liver within CL and ML region specifically but not PV region. N = 5 mice for each group. (**j**) Dot plot graph shows BrdU positive hepatocytes with the bigger nuclear area as compared with control groups. Cell number (n) = 300 for each group. (**k**) Immunostaining images of liver sections illustrate the cell and nucleus size of hepatocytes from age-matched control and drugs-treated livers including ATFB1, $CCl_4$, and 45 kcal% HFD at the indicated times. One-way ANOVA with Bonferroni's post-test was used to (**c**) and (**f**); Two-way ANOVA with Bonferroni's post-test was applied to (**i**). Student's unpaired t-test with Welch correction was used in (**j**); Values represent the mean ± s.e.m, *$P < 0.05$, **$P < 0.01$, ***$P < 0.001$. n.s., not significant. Scale bars: 200 μm in (**e**) and (**h**), 100 μm in (**a**), (**c**), and (**k**). 50 μm in high power field of (**c**). 30 μm in high magnification of (**k**).

**Supplementary Figure 2**

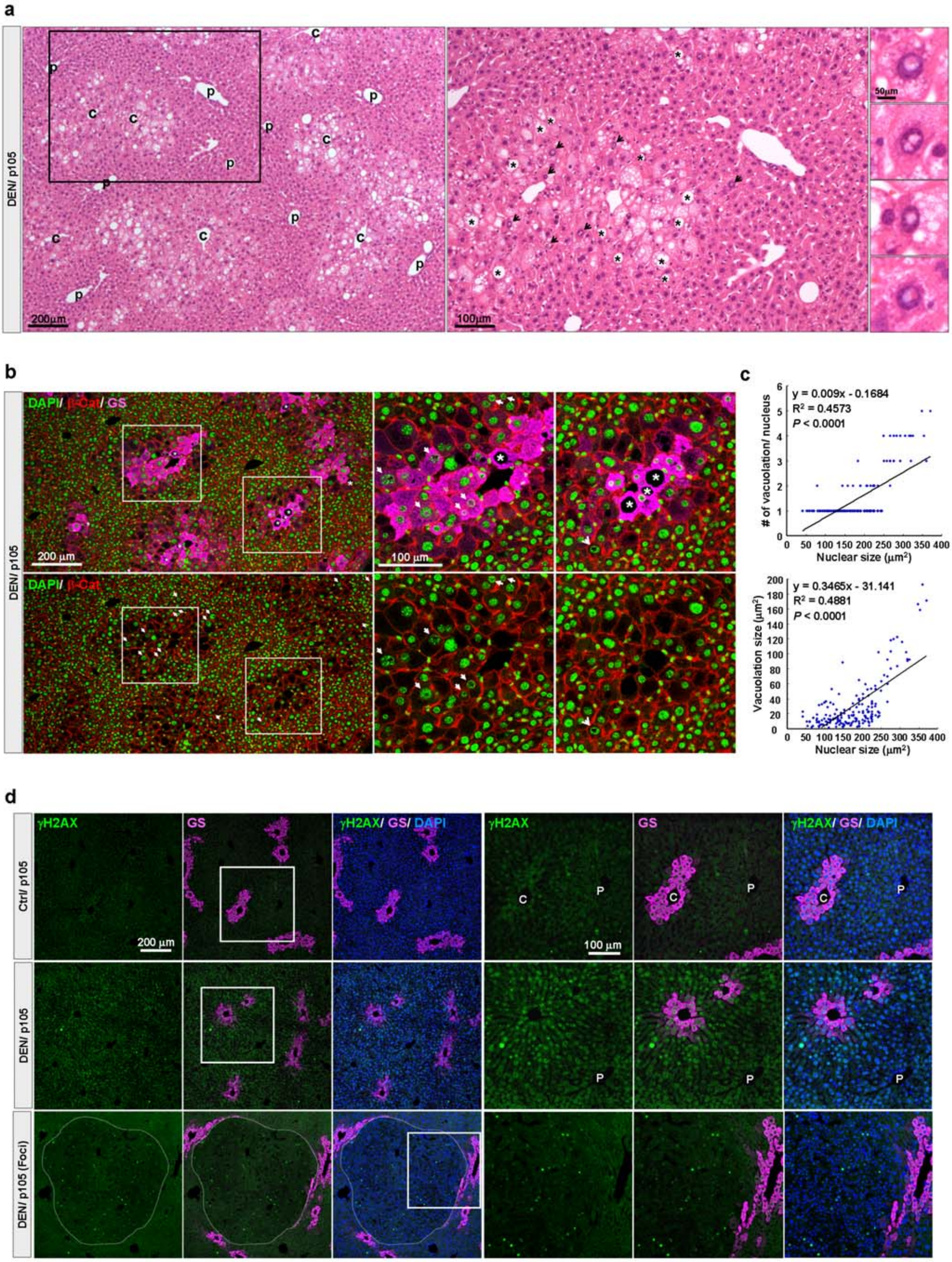

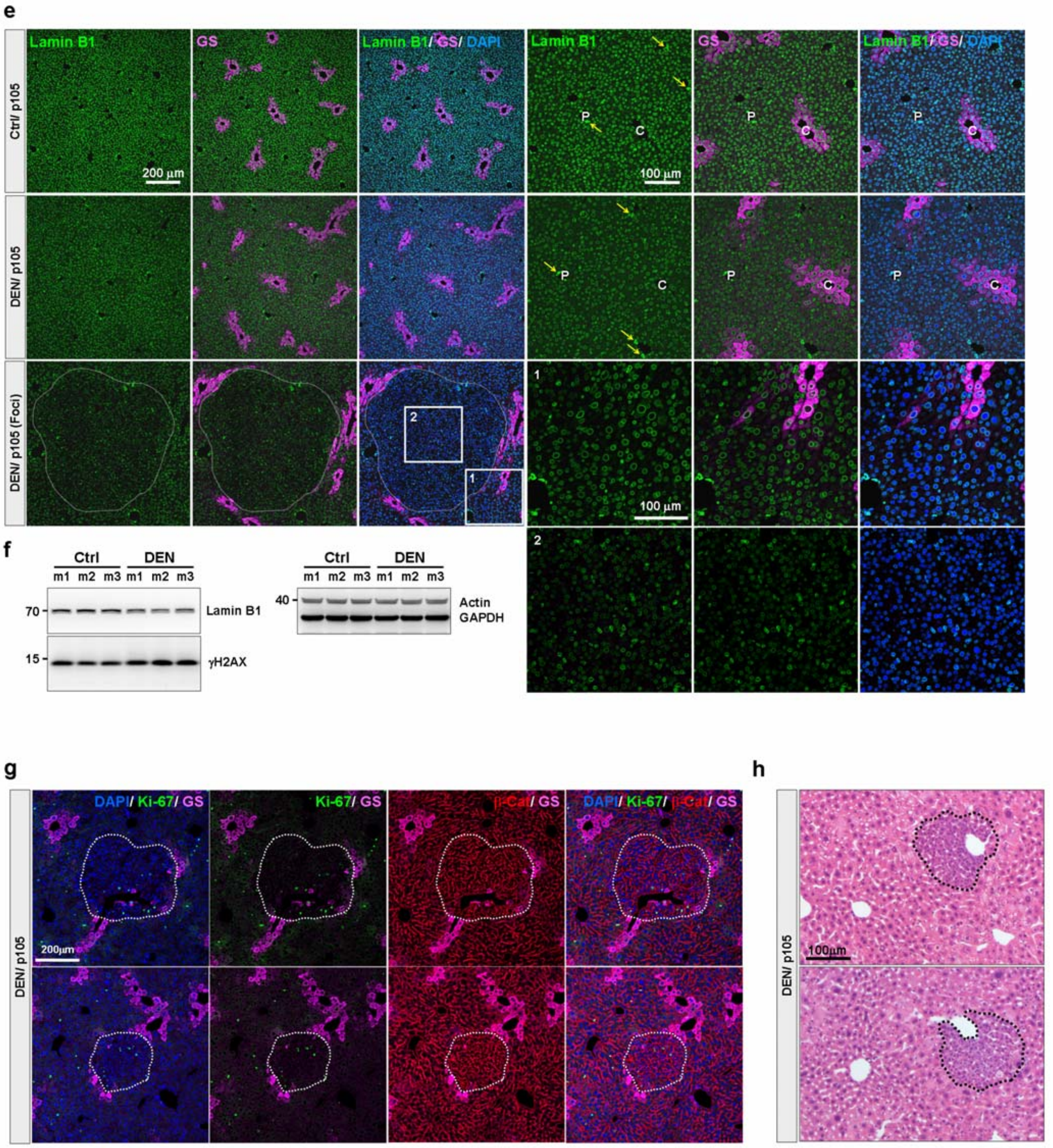
e
Ctrl/ p105
DEN/ p105
DEN/ p105 (Foci)
Lamin B1
GS
Lamin B1/ GS/ DAPI
200 µm
100 µm
P
C
1
2
f
Ctrl
DEN
m1 m2 m3 m1 m2 m3
70
Lamin B1
15
γH2AX
40
Actin
GAPDH
g
DEN/ p105
DAPI/ Ki-67/ GS
Ki-67/ GS
β-Cat/ GS
DAPI/ Ki-67/ β-Cat/ GS
200µm
h
100µm

**Supplementary Figure 2.** DEN treatment causes appearance of preneoplastic lesions within CL and ML region. (**a**) Images of 3 months DEN-treated liver with H&E staining show the nuclear (arrow) and cytoplasmic (asterisk) vacuolation dominantly within CL and ML region of liver. High magnification displays the detail structure of nuclear and cytoplasmic vacuolation, which are located nearby CL and ML region. (**b**) Nuclear and cellular morphology of 3 months DEN-treated livers were outlined by staining with DAPI (green) and $\beta$-catenin (red), and GS (magenta). The high magnification images show that nuclear (arrow) and cytoplasmic (asterisk) vacuolation are detected in GS-positive hepatocytes, and GS signal were observed within the vacuolation region of nucleus. (**c**) The correlation of the nuclear vacuolation number and size to nuclear size of hepatocytes was analyzed by linear regression. Each point represents an individual case from a hepatocyte. n > 170 hepatocytes with nuclear vacuolation from 5 mice. (**d**) The liver sections from control and 3 months DEN-treated mice were immunostained for $\gamma$H2AX (green) and GS (magenta). DNA content was visualized by DAPI signal (blue). Highly expressed $\gamma$H2AX was observed in hepatocytes adjacent to CL region and preneoplastic foci (dashed line). (**e**) Representative immunostaning images indicate the expression of Lamin B1 (green) in the liver. Nuclei and CL region were outlined by DAPI (Blue) and GS signal (magenta), respectively. Three months DEN-treated hepatocytes showed lower expression of Lamin B1 than those in control; and the lowest Lamin B1 signal was observed within preneoplastic foci. Note that Lamin B1 signal showed similar expression level in portal triad cells (yellow arrow) in all groups. P indicates portal vein region. (**f**) Protein level of $\gamma$H2AX and Lamin B1 in control and DEN-treated livers were examined by immunoblotting. (**g**-**h**) Immunohistochemistry (**g**) and H&E staining (**h**) show the dominant distribution of preneoplastic foci (dashed circles) nearby GS positive hepatocytes and CL region of the liver. Ki-67 positive signal was enriched within preneoplastic foci. For immunohistochemistry, liver slices were co-stained with DAPI (blue), Ki-67 (green), $\beta$-catenin (red), and GS (magenta). Linear regression was used to (**c**). Values represent the mean $\pm$ s.e.m, *$P < 0.05$, **$P < 0.01$, ***$P < 0.001$. Scale bars: 200 $\mu$m in low magnification of (**a**), (**b**), (**d**), (**e**), and (**g**), 100 μm in high magnification of (**a**), (**b**), (**d**), (**e**), and (**h**), and 50 μm in nuclear vacuolation images of (**a**).

**Supplementary Figure 3**

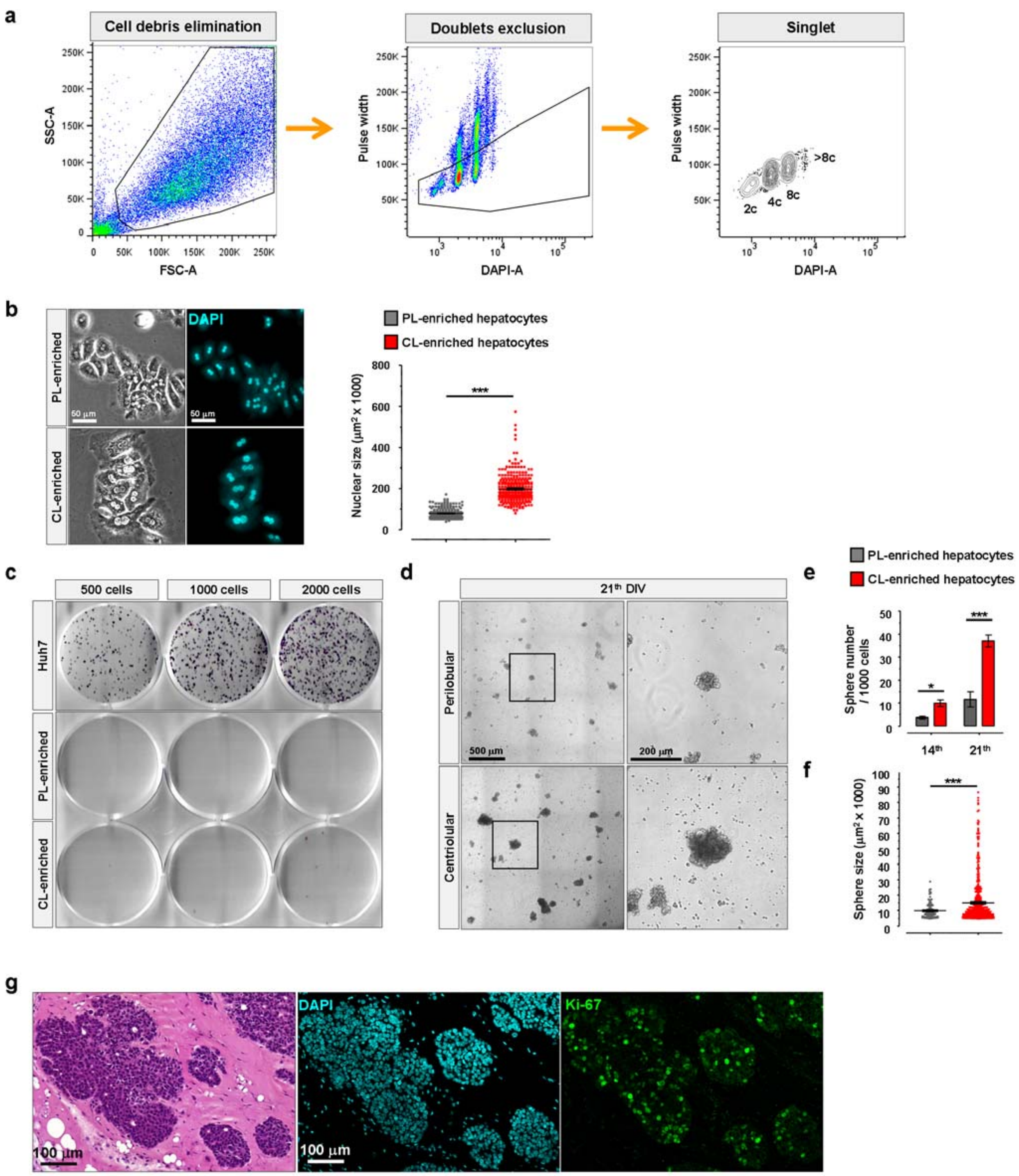

**Supplementary Figure 3.** CL-enriched hepatocytes show higher tumor stem cell characteristic. (**a**) Schematic illustration of gating strategy for the quantification of hepatocyte DNA content by flow cytometry after isolation through digitonin-collagenase infusion protocol from mouse livers. Abbreviation: FSC-A, forward scatter area; SSC-A, side scatter area. (**b**) Representative images display the cellular morphology and nucleus size of CL- or PL-enriched hepatocytes after 36h of seeding. Nuclear size was outlined by DAPI staining, and the quantitative data indicates that CV-enriched hepatocytes display larger nucleus size compared with PV-enriched hepatocytes. n = 300 hepatocytes/group. (**c**) Representative images show the colony formation assay with indicated cultured cell numbers in 6-well plate. Cells were fixed at 21$^{th}$ DIV and stained with crystal violet for colony numbers analysis, and Huh7 cells are the positive control. (**d**) Example of images show the tumorsphere formation of CL- and PL-enriched hepatocytes at 21$^{th}$ DIV. (**e-f**) The quantitative data indicates that tumorspheres derived from CL-enriched hepatocytes display higher numbers and larger size compared to PL-enriched group. (**g**) Representative immunohistochemistry and H&E stained images display the tumor cells derived from CL-enriched hepatocytes after three months of subcutaneous transplantation in NSG™ mice. Ki-67 signal positive cells (green) were enriched within the tumor nodule. DNA content was counter stained with DAPI (cyan). Student's unpaired t-test with Welch correction was used in (**b**) and (**f**); Two-way ANOVA with Bonferroni's post-test was applied to sphere numbers of (**e**). Values represent the mean ± s.e.m, *$P$ < 0.05, **$P$ < 0.01, ***$P$ < 0.001. Scale bars: 500 μm in high magnification of (**d**), 200 μm in low magnification of (**d**), 100 μm in (**g**), and 50 μm in (**b**).

#### Supplementary Figure 4

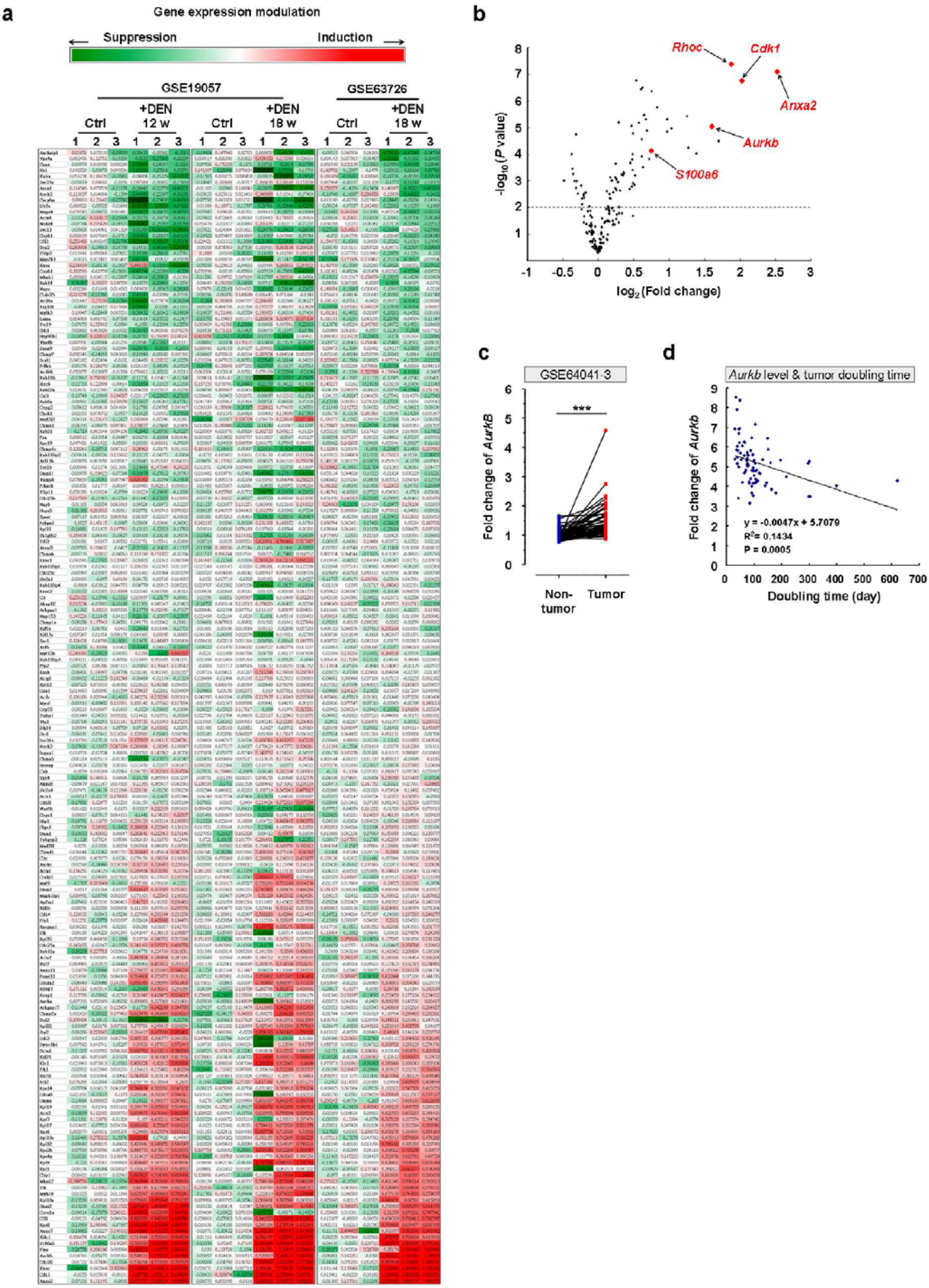

**Supplementary Figure 4.** *Aurkb* is a target for hyperpolyploidization of hepatocytes. (**a**) Heatmap of relative expression of cytokinesis genes up- or down-regulated by DEN in the liver. Cytokinesis genes were characterized and identified by previous studies[33-35], hepatic gene expression changes were obtained from two GEO datasets (GSE19057 and GSE63726)[36-37]. (**b**) Scatter plot graph with logarithmic and *P*-value axis shows the top five cytokinesis genes that are upregulated significantly in two GEO datasets. (**c**) Human *Aurkb* expression in matched normal and HCC biopsies, hepatic gene expression changes were obtained from two GEO datasets. Each pair data indicates one patient. N = 60 patients with HCC. (**d**) The scatter plot graph shows the linear regression analysis of the correlation between *Aurkb* expression and tumor doubling time of patient. Each point represents an individual case from a patient. N = 81 patients with HCC. Two-tailed Student's paired t-test was used in (**c**). Linear regression was conducted to (**d**). Values represent the mean ± s.e.m, $^{*}P < 0.05$, $^{**}P < 0.01$, $^{***}P < 0.001$.

#### Supplementary Figure 5

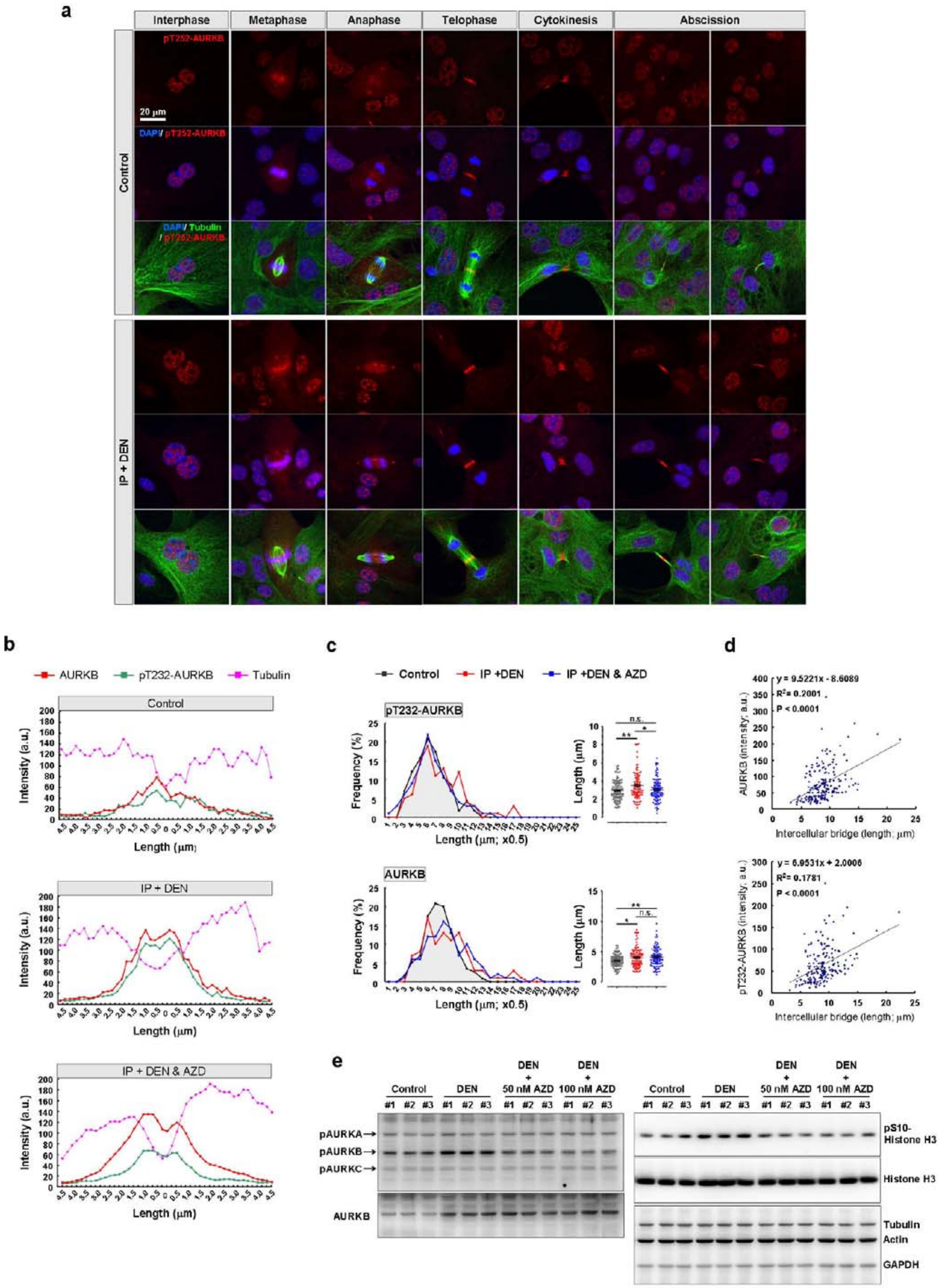

**Supplementary Figure 5.** Enrichment of pT232-AURKB at the midbody after DEN treatment. (**a**) Example of images show the immunofluorescence of pT232-AURKB (red) at different stage of cell cycle. Higher expression of pT232-AURKB was detected in hepatocytes isolated from liver with DEN treatment. Similar subcellular distribution of pT232-AURKB was observed between control and DEN-treated hepatocytes through whole cell cycle stages. Notable, highly enriched pT232-AURKB was expressed at the midbody during abscission stage in DEN-treated hepatocytes. The morphology of nuclei and intercellular bridges were outline by counterstaining with tubulin (green) and DAPI (blue) respectively. (**b**) Example of line-scan analysis of pT232-AURKB and AURKB intensity at the midbody (tubulin signal) from indicated cultured hepatocytes in Fig. 4h. (**c**) Frequency distribution and dot plot graphs show the length of AURKB and pT232-AURKB signal along the intercellular bridge. Longer AURKB and pT232-AURKB signal were detected in DEN-treated dividing hepatocytes compare to control group, that is rescued by 50 nM AZD1152 treatment. n > 100 dividing hepatocytes at abscission stage per group from three independent experiments. (**d**) Linear regression analysis shows the correlation between the expression level of AURKB and pT232-AURKB at the midbody and intercellular bridge length. n > 100 dividing hepatocytes at abscission stage from three independent experiments. (**e**) Immunoblots display that phosphorylation of AURKB at Thr232 residue and the downstream target signal of AURKB were inhibited by AZD1152 treatment. One-way ANOVA with Bonferroni's post-test was applied to (**c**). Linear regression was conducted to (**d**). Values represent the mean $\pm$ s.e.m, *$P$ < 0.05, **$P$ < 0.01, ***$P$ < 0.001. n.s., not significant. Scale bars: 20 $\mu$m in (**a**).

#### Supplementary Figure 6

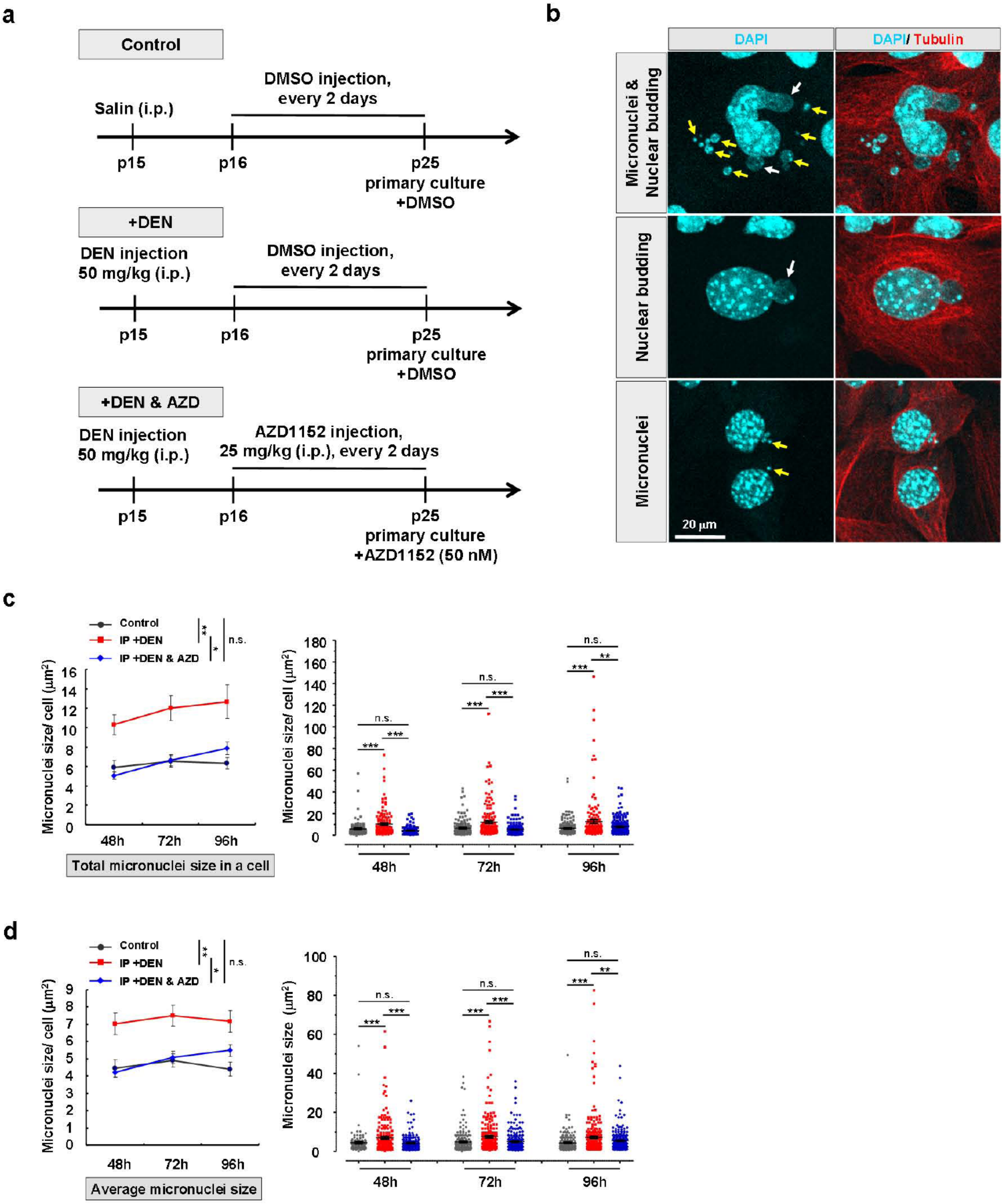

**e**

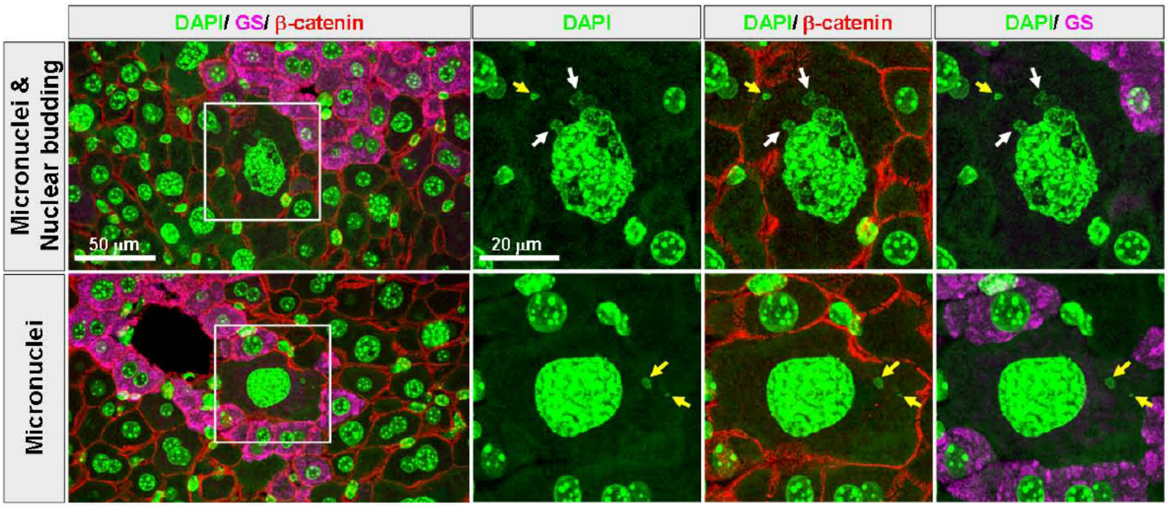

**f**

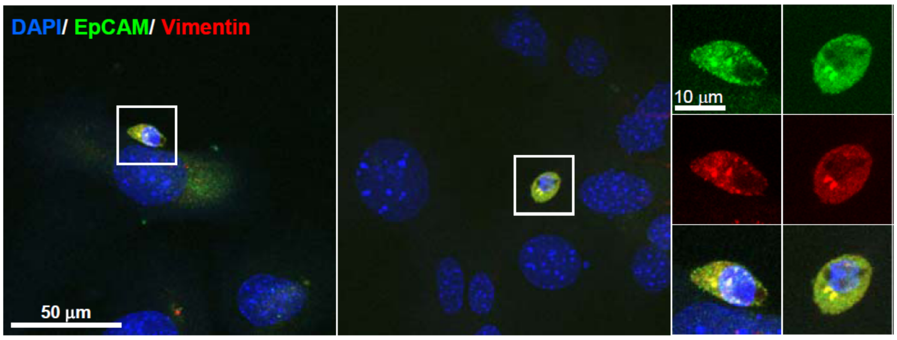

**g**

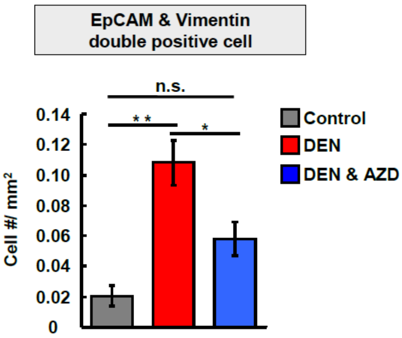

**Supplementary Figure 6.** Analysis the properties of nuclear budding and micronuclei in drugs treated cultured hepatocytes. (**a**) Schematic illustration of the hepatocyte primary culture and drugs administration protocol. Mice were injected with saline or DEN at p15 and sacrifice at p25. One day after DEN injection, AZD1152 (25 mg/kg) was applied to DEN-treated mice with every two days, while control and DEN only mice were treated with the same concentration of DMSO in saline. (**b**) Representative images show the typical morphology of micronuclei (yellow arrow) and nuclear budding (white arrow) in hepatocytes with DEN treatment. A variety of sizes and shapes of micronuclei and nuclear budding were observed. Micronuclei and nuclear budding have been defined as structures that are responsible for the expulsion of undesirable DNA content. Nuclear budding had a connection with the main nucleus, some were circular and positioned close to the nucleus, while some resided in the cytoplasm separated from the nucleus called micronuclei. (**c** and **a**) Total and average micronuclei size in a cell were analyzed showing a significant increase in

DEN-treated hepatocyte at indicated time points. Note that the size was dramatically reduced after AZD1152 treatment in DEN-treated group. n > 100 cells with micronuclei from three independent experiments. (**e**) Example of liver slice images displays micronuclei (yellow arrow) and nuclear budding (white arrow) within hyperpolyploid hepatocytes in the liver with 3 months of DEN treatment. (**f**-**g**) Immunocytochemistry shows example images of double staining cancer stem cell markers, EpCAM and Vimentin, in cultured hepatocytes isolated from DEN-injected liver after one week culture. Highly expression of both EpCAM and Vimentin was observed in cells with smaller nuclei which is adjacent to the hepatocytes with bigger nucleus size. Significantly increased the numbers of EpCAM and Vimentin double positive cell in cultured hepatocytes isolated from the DEN-treated liver. AZD treatment dramatically reduced the number of EpCAM and Vimentin double positive cell in DEN-treated hepatocytes. Two-way ANOVA with Bonferroni's post-test was applied to line graphs of (**c**), and (**d**); One-way ANOVA with Bonferroni's post-test was used to dot plot graphs of (**c**), (**d**), and (**g**). Values represent the mean ± s.e.m, *$P$ < 0.05, **$P$ < 0.01, ***$P$ < 0.001. n.s., not significant. Scale bars: 50 μm in low magnification of (**e**) and (**f**), 20 μm in (**b**) and high magnification of (**e**), 10 μm in high magnification of (**f**).

**Supplementary Figure 7**

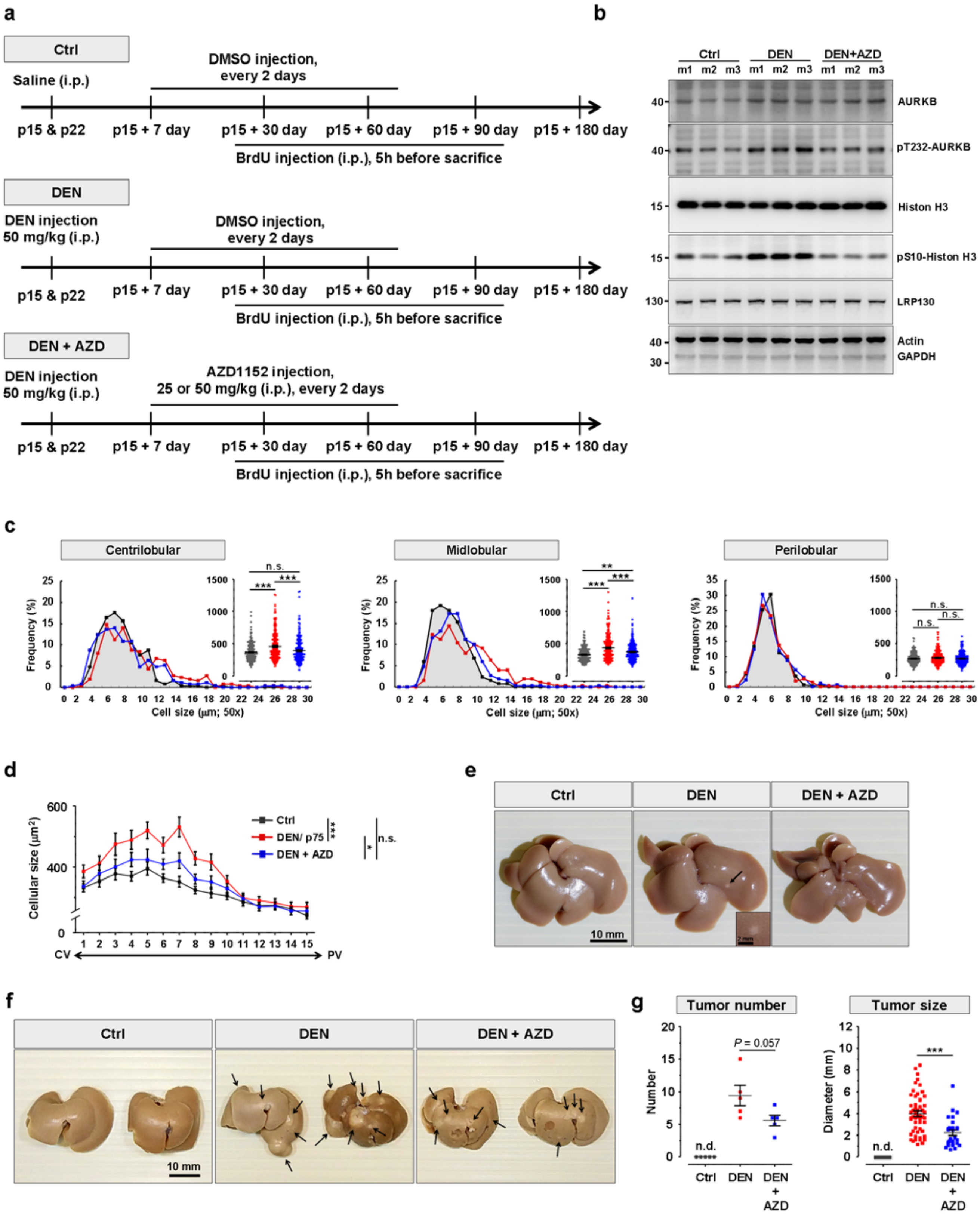

**Supplementary Figure 7.** Inhibition of AURKB activity reduces cell size of hepatocytes in DEN-treated liver. (**a**) Schematic illustration of drugs administration protocol in mice. Mice were injected with saline or DEN twice at p15 and p22. One week after DEN injection, AZD1152 was applied to DEN-treated mice every two days, while control and DEN only group were treated with the same concentration of DMSO in saline. (**b**) Western blots show the significant decrease of pT232-AURKB in DEN-treated liver after one month of AZD1152 (25 mg/kg) injection. Phosphorylation of histone H3 at Ser10 residue, downstream signal of AURKB, were reduced in one month of AZD1152 treated livers. Protein loading equivalence was controlled by b-actin, GAPDH, and LRP130 levels, which stands for the internal control at different molecule size. (**c**) Frequency distribution and dot plot graphs show the quantitative data of cell size of hepatocytes within specific liver lobule in indicated mice. N = 5 mice per group, and n = 750 hepatocytes per group. (**d**) Frequency distribution of cellular size of hepatocytes along CV-PV axis in indicated mice. N = 5 mice and n = 750 hepatocytes for each group. (**e**) Example of liver images display the global morphology livers with or without drugs treatment. The general morphology shows no significant change between control and DEN-treated liver, although very few tumor nodules were found on the surface of DEN-treated liver infrequently. No tumor nodules were discovered on the surface of DEN-treated liver followed AZD1152 (25 mg/kg) treatment. (**f**) Representative livers from control (Ctrl), DEN-treatment (DEN), DEN-treatment followed by 2 month AZD1152 (DEN+AZD) injection 6 month after DEN-injection. (**g**) Quantitative data show the tumor diameter and tumor numbers in Fig. 7f. N = 5 mice/group. One-way ANOVA with Bonferroni's post-test was used to (**c**) and (**g**); Two-way ANOVA with Bonferroni's post-test was applied to (**d**). Values represent the mean ± s.e.m, **P* < 0.05, ***P* < 0.01, ****P* < 0.001. n.s., not significant; n.d., not detectable. Scale bars: 10 mm in (**e** and **f**).

**Supplementary Table 1**

**Table 1.** Primer pairs utilized in qPCR.

| Oligo title | Sequence (5' to 3') | UPL probe # |
|---|---|---|
| Anxa2-f | ggaaatatggcaagtccctgt | #42 |
| Anxa2-r | tctggtagtcacccttggtgt | |
| Aurkb-f | attgcagactttggctggtc | #69 |
| Aurkb-r | aatcatctctgggggcagat | |
| Cdk1-f | gaacttcgacatccaaatatagtcag | #64 |
| Cdk1-r | ccatggacaggaactcaaaga | |
| Rhoc-f | aaggacctgaggcaagatga | #92 |
| Rhoc-r | aaggcactgatcctgtttgc | |
| S100a6-f | aggaaggtgacaagcacacc | #17 |
| S100a6-r | agcatcctgcagcttgga | |
| GS-f | gagcccaagtgtgtggaag | #58 |
| GS-r | aaggggtctcgaaacatgg | |
| Axin2-f | ttattgctactccaaatgcaaaag | #50 |
| Axin2-r | ttttggcaaggtaccacctc | |
| Rhbg-f | tcacactggtgtttgcctct | #18 |
| Rhbg-r | gaagcattgggagtctggag | |
| Oat-f | taacgatctgcccgcact | #22 |
| Oat-r | aacgataacgcctgcttcac | |
| Lect2-f | gcaccattcactgggaagata | #02 |
| Lect2-r | tgtagaaaattttgacacaaaaacct | |
| Pck1-f | ggagtacccattgagggtatcat | #49 |
| Pck1-r | gctgagggcttcatagacaag | |
| Gls2-f | tgacttctcgggccagttt | #88 |
| Gls2-r | gcccatgacattgggtaca | |
| Arg1-f | cctgaaggaactgaaaggaaag | #02 |
| Arg1-r | ttggcagatatgcagggagt | |
| Cps1-f | ccctctgactatgttgccatt | #72 |
| Cps1-r | gggtcagcatctctcagtcg | |
| Cyp2f2-f | aaataccccaaagtgcaagc | #11 |
| Cyp2f2-r | tgcatctgtgtaaggcatgg | |
| Actb-f | ctaaggccaaccgtgaaaag | #64 |
| Actb-r | accagaggcatacagggaca | |
| Gapdh-f | gggttcctataaatacggactgc | #52 |
| Gapdh-r | ccattttgtctacgggacga | |
